# Patient derived organoids reveal that PI3K/AKT signalling is an escape pathway for radioresistance and a target for therapy in rectal cancer

**DOI:** 10.1101/2021.08.31.458326

**Authors:** Kasun Wanigasooriya, Joao D. Barros-Silva, Louise Tee, Mohammed E. El-Asrag, Agata Stodolna, Oliver J. Pickles, Joanne Stockton, Claire Bryer, Rachel Hoare, Celina Whalley, Robert Tyler, Tortieju Sillo, Christopher Yau, Tariq Ismail, Andrew D. Beggs

## Abstract

Partial or total resistance to preoperative chemoradiotherapy occurs in more than half of locally advanced rectal cancer patients. Several novel or repurposed drugs have been trialled to improve cancer cell sensitivity to radiotherapy, with limited success. To understand the mechanisms underlying this resistance and target them effectively, we initially compared treatment-naive transcriptomes of radiation-resistant and radiation-sensitive patient-derived organoids (PDO) to identify biological pathways involved in radiation resistance. Pathway analysis revealed that PI3K/AKT/mTOR and epithelial mesenchymal transition pathway genes were upregulated in radioresistant PDOs. Moreover, single-cell sequencing of pre & post-irradiation PDOs showed mTORC1 upregulation, which was confirmed by a genome-wide CRSIPR-Cas9 knockout screen using irradiated colorectal cancer (CRC) cell lines. Based on these findings, we evaluated cancer cell viability *in vitro* when treated with radiation in combination with dual PI3K/mTOR inhibitors apitolisib or dactolisib. Significant AKT phosphorylation was detected in HCT116 cells two hours post-irradiation (*p*=0.027). Dual PI3K/mTOR inhibitors radiosensitised HCT116 and radiation-resistant PDO lines. The PI3K/AKT/mTOR pathway upregulation contributes to radioresistance and its pharmacological inhibition leads to significant radiosensitisation in an organoid model of CRC and is a target for clinical trials.

## 1. Introduction

Neoadjuvant chemoradiotherapy (CRT) followed by surgical resection is the standard of care for patients with locally advanced rectal cancer (LARC).^1^ CRT downstages the disease, increases rates of sphincter sparing surgery and reduces the risk of local recurrence.^1–5^ Some patients may be treated with a short course of neoadjuvant radiotherapy (SCRT).^6^ Approximately 10-30% of patients receiving CRT and 2-6% receiving SCRT demonstrate a complete pathological response (pCR), where no residual tumour cells are seen on histopathological assessment of post-resection surgical specimens.^7–9^ Conversely, around a third of LARC patients demonstrate no response to CRT.^10^ Various clinicopathological features,^11–18^ biomarkers: the tumour microenvironment,^19^ microbiome,^20, 21^ and genomic markers^22–25^ have been linked to pCR. However, a definitive molecular mechanism underlying CRT resistance in rectal cancer cells remains unclear. Understanding the pathways that drive CRT resistance is essential to developing tools to predict patient response and improve treatment response.^25^

The impact of tumour heterogeneity and acquired resistance to therapies, and inter-patient variability to treatment response, are now widely acknowledged.^26^ Patient-derived organoids (PDO) can be used as preclinical research models that resemble the genomic heterogeneity observed in patients, whilst allowing the reliability and flexibility required of *in vitro* assays, and have been recently used to study rectal cancer.^27, 28^ PDOs maintain cellular heterogeneity and genetic stability after multiple passages and respond similarly to treatment both in vivo and in vitro, rendering them a powerful tool in cancer research.^26^ PDOs have been developed from primary RC biopsies and surgical resection specimens recapitulating the genetic diversity and treatment response within RCs.^27, 28^ Ganesh et al. demonstrated that RC organoids mirror clinical responses of individual patients to chemoradiotherapy.^27^ Yao et al. observed a broad range of intrinsic PDO responses to conventional chemoradiation.^28^ Theses studies suggest PDOs can serve as tumour avatars which predict RC response to various treatments.

In the present study we utilised PDO models to investigate the mechanisms of resistance to radiotherapy and test drugs that may improve radiotherapy response in RC. Comparative transcriptome analysis of radiosensitive and radioresistant PDOs revealed novel genes and pathways involved in radioresistance. Single cell sequencing of PDOs and genome-wide clustered regularly interspaced short palindromic repeats (CRISPR)–associated nuclease 9 (Cas9) knockout screens using CRC cell lines were performed to validate findings. Finally, we performed *in vitro* assays combining dual PI3K and mTOR inhibitors with the current standard chemotherapy agent (5FU) to evaluate how it affected response to radiation.

## 2. Results

### 2.1 Patient-derived organoids (PDO) derivation and characterisation

The first six out of a total stably derived 16 PDO lines derived in our laboratory were used in this study. To confirm tumour status (pan-Cytokeratin) and colorectal origin (CDX2) expression in PDOs was assessed using immunohistochemistry.

Immunostaining revealed pan-Cytokeratin expression in six PDO lines matching the primary tumours’ expression (Figure 1A). Two PDO lines (653 and 557) showed no staining for the intestinal-specific transcription factor CDX2, matching their primary tumours (Figure 1A; Supplementary Figure 1). CDX2 is an intestinal specific transcription factor and approximately 20% of CRC do not express CDX2.^29, 30^ Targeted DNA sequencing of PDO lines revealed mutations in 20 of the 30 genes included in the panel, with *APC* being mutated in all six PDO lines (Figure 1B; Supplementary Table 2). To evaluate PDO sensitivity to radiotherapy, each line was irradiated with daily fractions – ranging from 2 Gy to 40 Gy – over 5 days, after which cell viability was assessed (Figure 2A). Relative organoid cell viability decreased by 90% in four PDO lines (884, 064, 389 and 411) when treated with a total of 25 Gy or higher, whereas for lines 653 and 557 there was a 40% reduction (Figure 2B). The former group was deemed radiosensitive and the latter radioresistant. Targeted DNA sequencing revealed *KRAS* mutations (*G12D and G13R)* in the two radioresistant PDO lines, with line 653 additionally harbouring a *TP53* deletion *(P191del)* and a missense *FBXW7* mutation *(R465H)*. The 557 PDO line contained a *MSH6* frameshift mutation *(F1104fs)*, and the parent tumour also displayed deficient mismatch repair (dMMR) phenotype on immunohistochemistry. This line also harboured a pathological missense *PIK3CA* mutation (*R88Q)*. CDX2 immunostaining was detected in radiosensitive lines but absent in radioresistant PDOs (Figure 1A).

**Figure 1:**
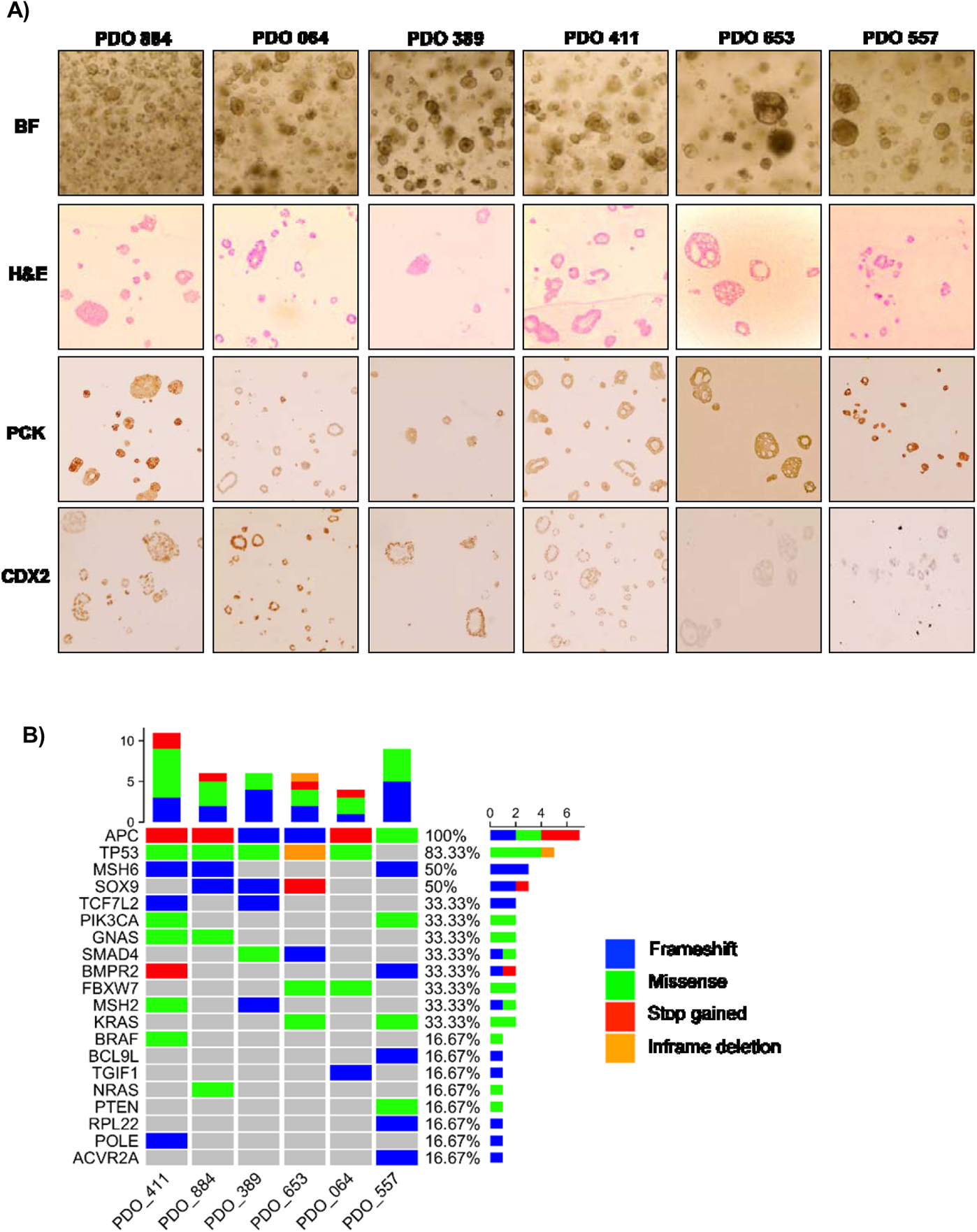
Optical microscopy, immunohistochemical and genomic characterisation of six PDO lines. A) Bright-field (BF) images and H&E staining of cultured PDOs. Immunohistochemistry staining of pan-Cytokeratin and CDX2 was performed to confirm the PDOs match the corresponding primary tumours. B) Targeted sequencing was performed using a panel of 30 known CRC genes (see methods for full list). The heatmap summarizes the mutational landscape identified in the PDO lines.

**Figure 2:**
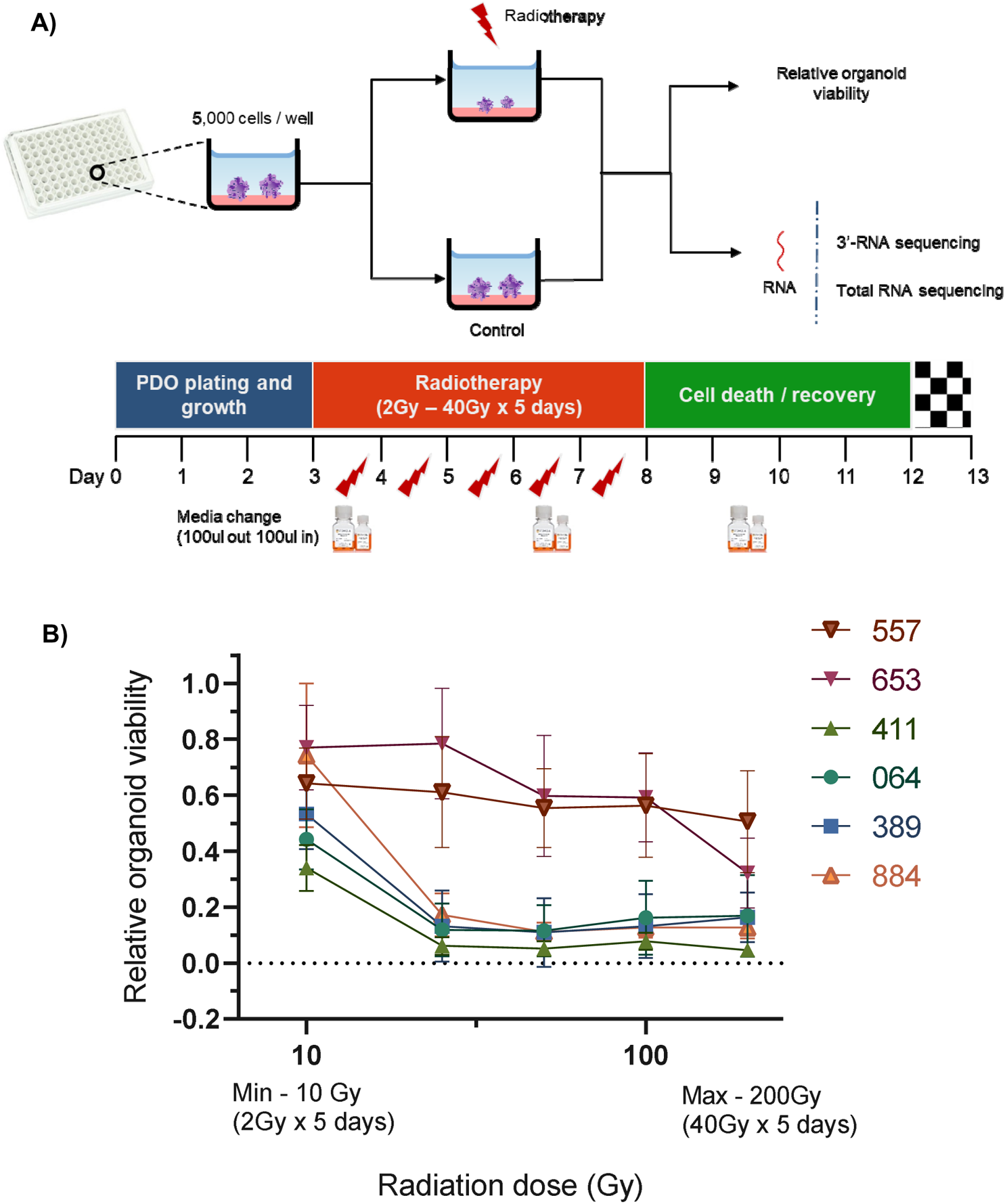
Radiotherapy response of the six PDO lines. A) The six PDO lines were plated at an estimated 5000cells/well of organoids in 96-well cell culture plates and were treated with different radiotherapy doses ranging from 2 Gy x 5 days to 40 Gy x 5 days. Relative PDO viability was assessed at the end of experiment through chemiluminescence detection using CellTiter-Glo® 3D. B) Combined normalised results from three independent experiments conducted for each PDO line has been displayed in this figure. Copyright of STEMCELL Technologies (Canada). Image used with permission. Source: https://www.stemcell.com/cell-separation/immunomagnetic

### 2.2 Differential transcriptional profiles of radioresistant organoids

Principal component analysis of the pre-treatment PDO transcriptomes grouped the samples two main clusters (Figure 3A) matching the radiation-response status we defined experimentally (Figure 2A), suggesting distinctive gene expression underlying radiosensitivity. The expression profiles of the two radioresistant PDO lines were more closely clustered compared to the radiosensitive PDO lines, identified using our earlier experiments (Figure 3A). In total, we observed 9659 differentially expressed genes (fold change [FC] >= 1.5) between the radioresistant and radiosensitive PDO lines (False discovery rate [FDR]< 0.05). Of these, 7294 were upregulated in radioresistant PDO lines and the remainder were upregulated in the radiosensitive lines. The top five significantly upregulated genes in the radioresistant PDO lines included *SCARA3, CAV1, NTN1, UBASH3B* and *PLK2,* whereas *OLFM4, SLC39A5, ABCB1*, *TDGF1,* and *CYP2B6* were the top five differentially upregulated genes in a radiosensitive PDO lines (Figure 3B). To further characterize the transcriptional differences, we performed pathway enrichment analyses using the Hallmark, KEGG and Oncogenic Signatures gene-set collection of the Molecular Signatures Database (MSigDB V7.4).

**Figure 3:**
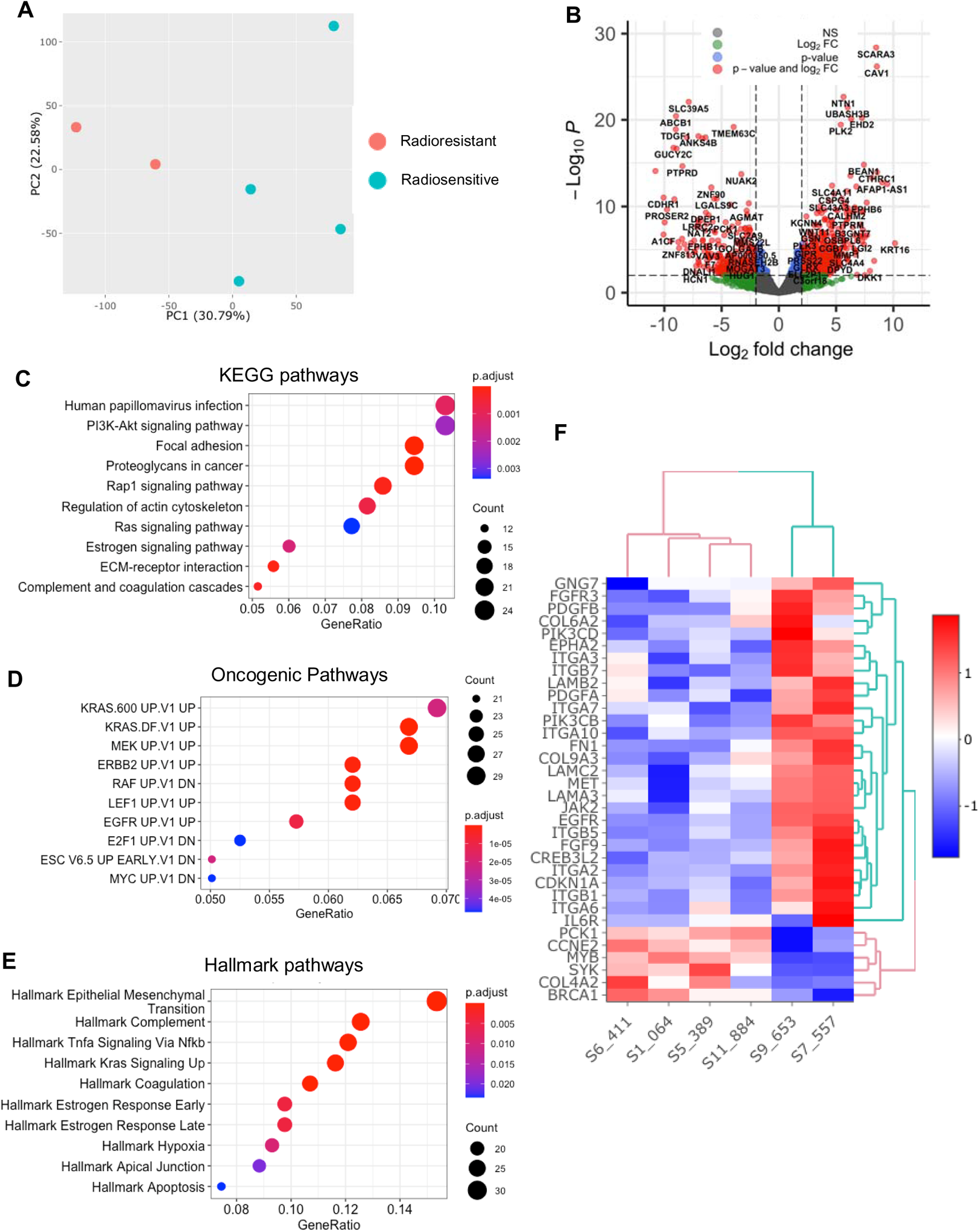
Gene expression at baseline of responding vs. non-responding PDO lines. A) Principal component analysis; B) Volcano plot demonstrating upregulated and downregulated genes in radioresistant PDOs; C-E) Gene set enrichment analysis using Kegg, Oncogenic Signatures and Hallmarks gene sets. F) Heatmap generated through hierarchical clustering demonstrating PI3K / AKT / mTOR pathway gene upregulation or downregulation amongst radiosensitive and radioresistant PDOs. (PC=principal component, FC=fold change)

KEGG enrichment analysis revealed several the Human papillomavirus infection pathway genes (24 of 233, *p*=0.001) and the PI3K/AKT signalling pathway genes (24 of 233, *p*=0.002) upregulated in the radioresistant PDOs (Figure 3C). The upregulated genes of the latter are represented in Figure 3F. Enrichment analysis using the MSigDB Oncogenic Signature gene set revealed that the upregulated genes in radioresistant PDO lines match a set of genes upregulated in *KRAS* mutant cells (Figure 3D), and Hallmark pathway analyses also revealed upregulation of Epithelial Mesenchymal Transition related genes (Figure 3D and 3E).

### 2.3 Genomic indicators of PI3K/AKT/mTOR pathway significance in radiation response

In order to understand the role of heterogeneity in response to radiation therapy, single cell sequencing was performed on three paired irradiated and control PDOs. A total of 51,000 cells with 24,011 gene features were identified, which on dimensionality reduction and manual annotation of clusters collapsed into 11 clusters that were further grouped in two larger clusters (Figure 4A and 4B). The first consisted of *CEACAM* expressing tumour cells, colonic stem cells and tumour cells with high epithelial markers (Figure 4B). The second cluster consisted of tumour cells with a small intestinal expression phenotype, dead and dying cells (expressing predominantly ribosomal genes), and tumour cells that overexpressed either *ELOB*, *YBX1* or *NEAT1* (Figure 4B). When the cells were separated by irradiated and non-irradiated groups, a large expansion in the cluster expressing colonic stem cells was identified. Pathway analysis by Seurat and PROGENy did not demonstrate any consistently dysregulated pathways (Figure 4C). Therefore, dimensionality reduction using Scalable Bayesian Boolean Tensor Factorisation (SBBTF) and pathway analysis was carried out, which showed significant upregulation of mTOR, DNA damage repair, mitochondrial translation and electron transport in irradiated organoids, and significant downregulation of Hedgehog signalling and the cell cycle (Figure 4D and 4E).

**Figure 4:**
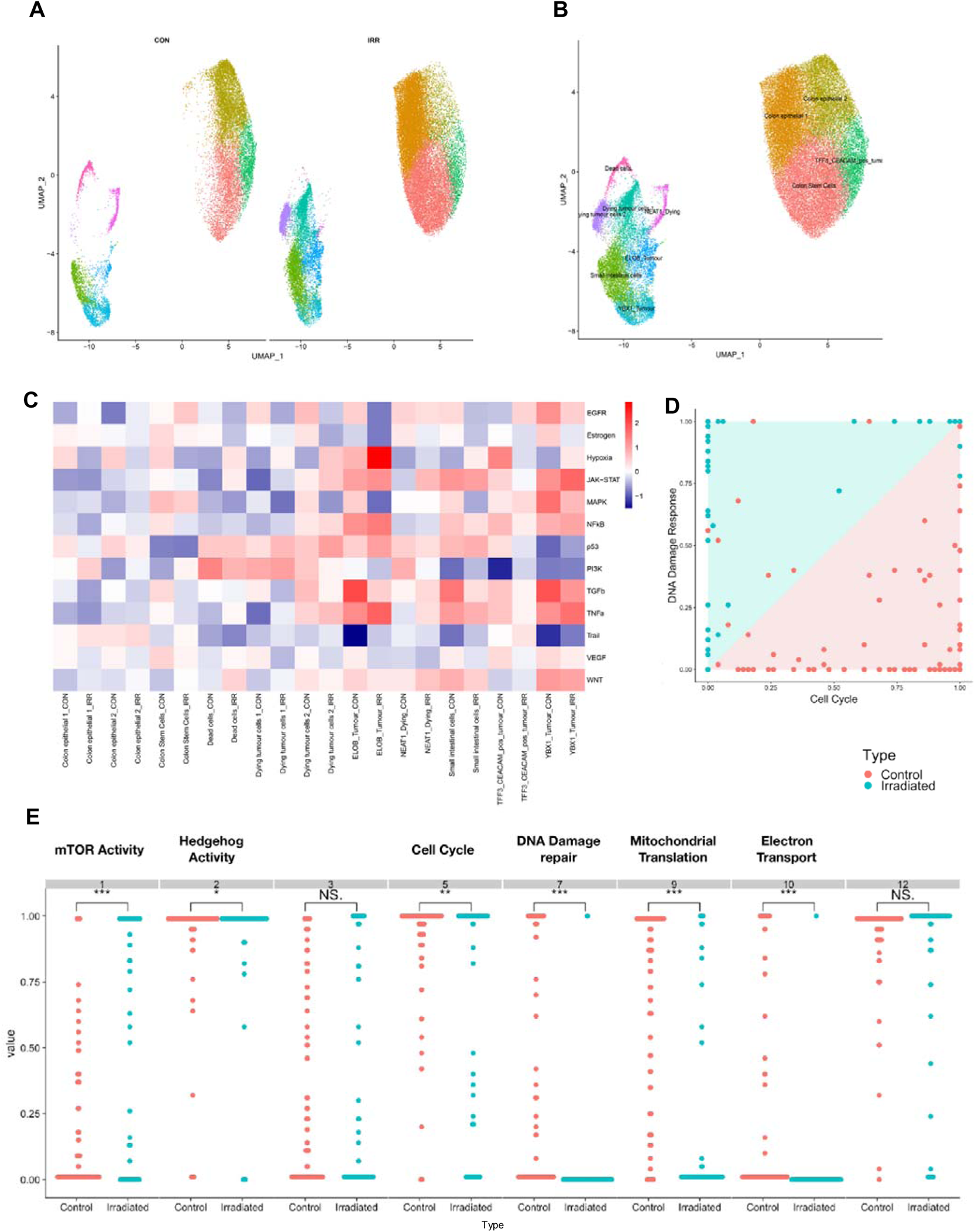
Single cell sequencing of irradiated and non-irradiated PDOs. A) and B) Dimensionality clustering and manual annotation of single cell sequencing data. C) Heatmap generated through hierarchical clustering after pathway analysis using Seurat and PROGENy. D) and E) Results from Scalable Bayesian Boolean Tensor Factorisation (SBBTF) and pathway analysis. Con=control; IRR=irradiated.

To validate this result, a genome-wide CRISPR-Cas9 knockout screen was performed on the C80 and HT55 CRC cell lines. Differential sgRNA data were processed using MaGECKFlute identifying positively selected genes under the selection pressures of 5 Gy/day irradiation over 5 days (Figure 5). The volcano plot in Figure 5A depicts significantly differentially expressed genes (*p*<0.05 and FC> 1) in viable cells following irradiation. In total, there were 26 genes that showed expression based on the mTOR signalling pathway gene list from the MSigDB. *PIK3CA*, *PIK3CB* and *EIF4B* were differentially upregulated in irradiated samples compared to control samples, and PTEN was downregulated in the irradiated cells (Figure 5A). Enrichment analysis using KEGG gene sets of the MSigDB revealed upregulated genes matched the PI3K / AKT / mTOR pathway gene set (Figure 5B). The upregulated genes also matched several pathways of the Oncogenic Signatures gene set collection (Figure 5C). These included the KRAS and WNT signalling.

**Figure 5:**
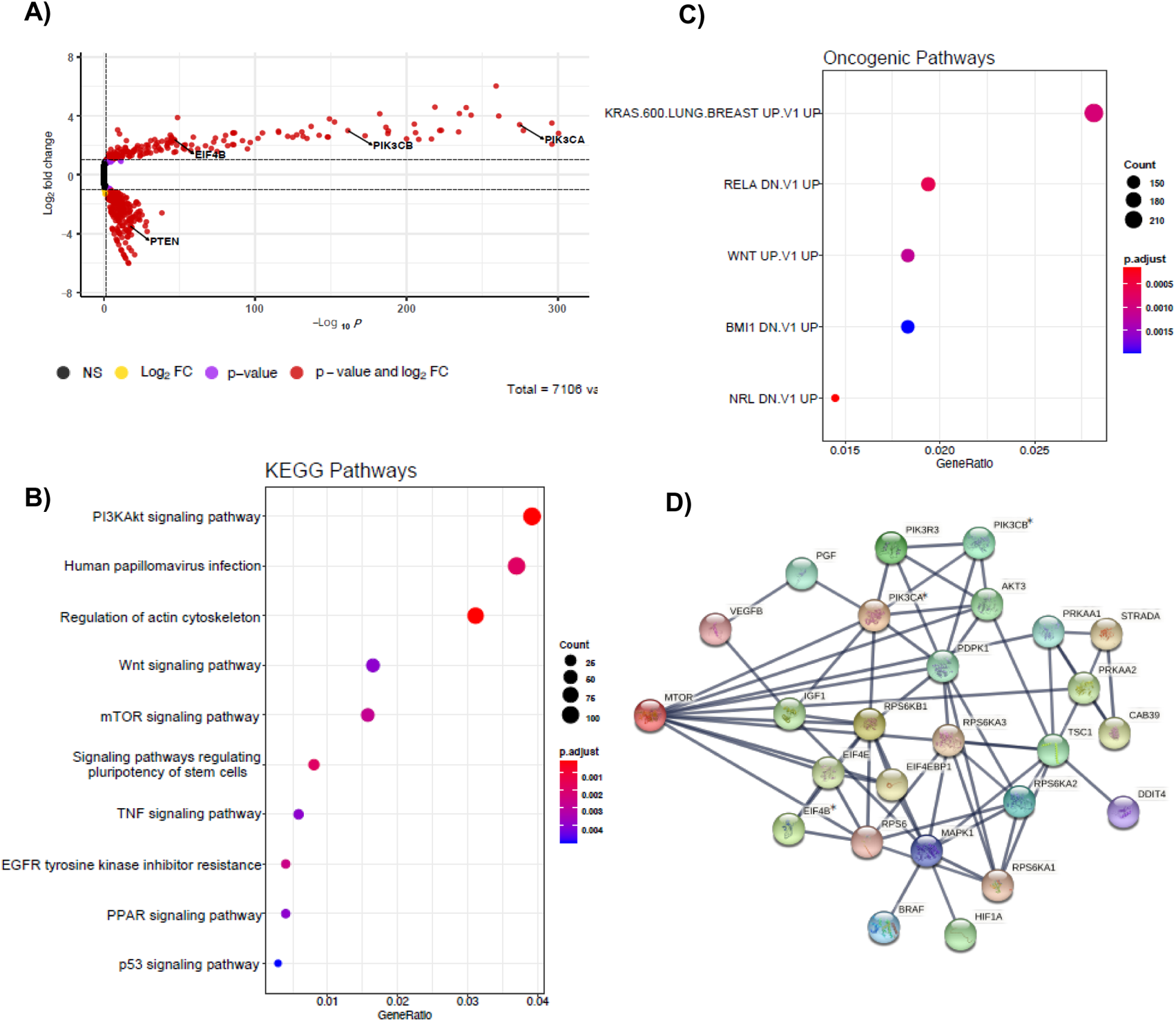
Genome-wide CRISPR screens identify mTOR signalling up-regulates pathway genes. A) Volcano plot depicting significantly differentially expressed genes (p < 0.05 and |fold change| > 1) in red. Model-based analysis of genome-wide CRISPR-Cas9 knockout on libraries A and B designed by Zhang lab 1 detected twenty-six expressed genes from mTOR signalling pathway gene list based on molecular signatures database (MSigDB) v7.4. B) Dot plot showing significantly over-represented KEGG pathway terms that were defined from MSigDB v7.4 using the 3,207 DEGs with p < 0.05. C) Dot plot showing significantly over-represented oncogenic pathway terms that were defined from MSigDB v7.4 using the 3,207 DEGs with p < 0.05. D) Network diagram representing the mTOR signalling events mediated by PIK3CA, PIK3CB and EIF4B using STRING 2, weighted correlation network analysis (WGCNA) with the Pearson’s correlation coefficients of every possible pair of genes were detected 12 modules by hierarchical clustering and subsequent module merging by eigengenes. Asterisks represent statistically significant levels of gene differences between control and irradiated samples (p< 0.05) calculated using Student’s t test. NS=non-significant; FC=fold change

Module eigengenes were calculated and hierarchically clustered to merge very closely correlated modules together, reducing the total number of modules. A total of 12 modules (groups of co-expressed genes) were detected by hierarchical clustering and subsequent module merging using eigengenes. mTOR signalling module were detected and visualized using STRING 4 (Figure 5D). Negative selection was observed in the entire PI3K/AKT/mTOR pathway, as well as the RAS/RAF/MEK/ERK signalling pathways, suggesting these pathways were involved with the cellular response to radiotherapy.

### 2.4 Dual PI3K/mTOR inhibitors improve CRC cell line response to radiation, however single agent mTOR inhibition does not

Following the findings of PIK3/AKT axis up-regulation in radioresistant PDO, we initially tested both sirolimus and everolimus effect on the HCT116 cell line combined with radiotherapy, finding no measurable difference (data not shown). We thenmade a decision to test dual inhibitors of PI3K/MTOR. HCT116 was treated *in vitro* using 5FU, apitolisib or dactolisib (Figure 6) with or without radiotherapy administered in 1 Gy fractions over 5 days. The dose-response curves in Figure 6 summarise the combined results from three independent experiments. Radiotherapy as a standalone treatment had minimal effect on endpoint cell viability (<10% reduction in cell viability). Increasing the doses of all three drugs had a significant impact on cell viability with or without radiotherapy (*p*<0.0001; Figure 6B). All three drugs showed radiosensitising effects. The IC50 for HCT116 treated with dual PI3K/mTOR inhibitors apitolisb and dactolisib were 0.6μM, 0.1μM without radiotherapy, and 0.3μM, 0.03μM in the presence of radiotherapy, respectively. Previous research has revealed maximum safe plasma concentrations (Cmax) of 0.320μM to 0.380μM, when 30mg or 40mg of oral apitolosib were administered to patients respectively.^31^

**Figure 6:**
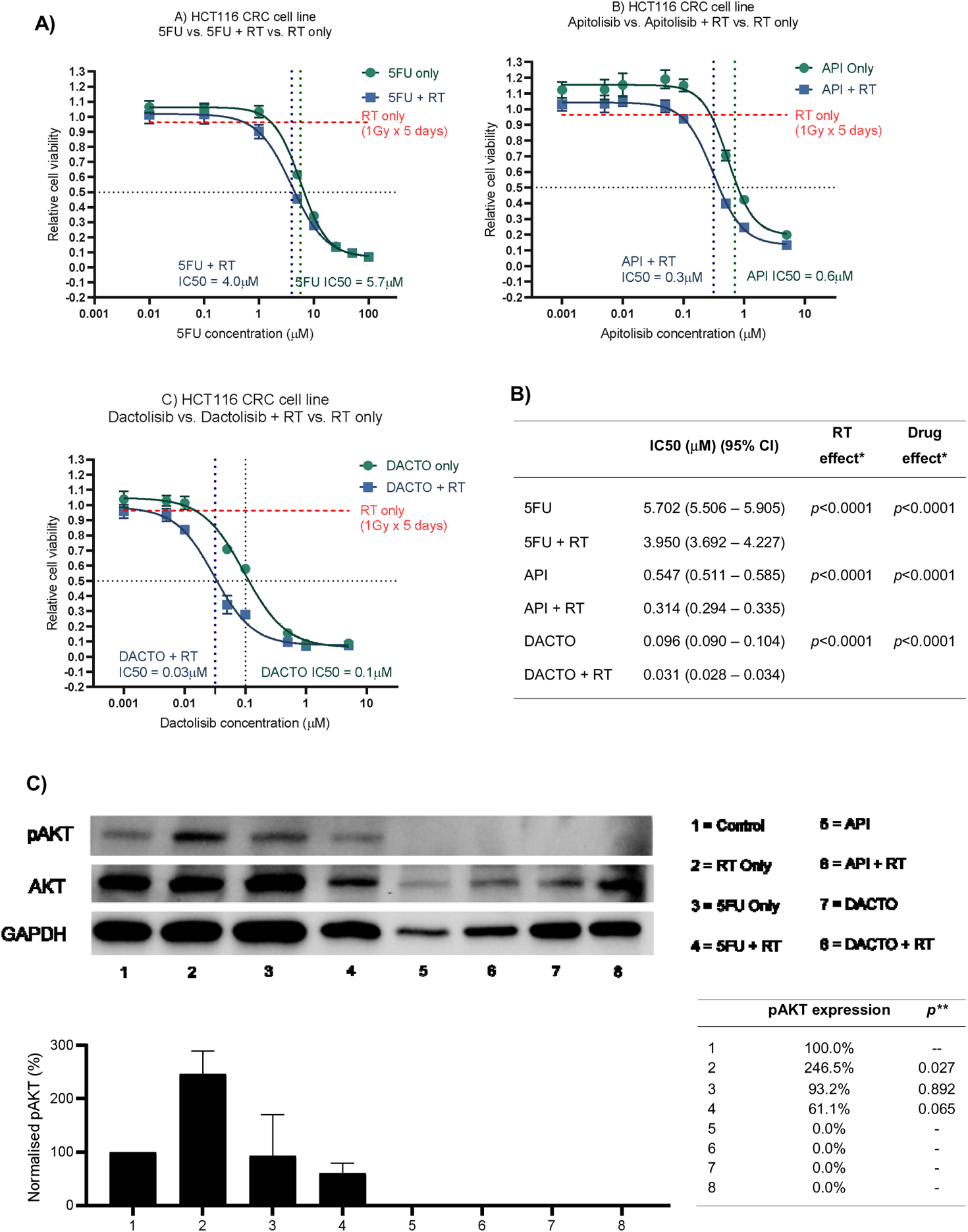
Radiosensitising HCT116 and its AKT phosphorylation status following drug and/or radiotherapy treatment. A) HCT116 was treated with eight different concentrations of 5-FU, Apitolisib or Dactolisib, with or without radiotherapy (1Gy fractions over 5 days). End-point cell viability was assessed using CellTiter-Glo 2.0 (Promega, USA) and chemiluminescence. The data is presented normalised to control (containing 0.1% DMSO). The x-axis represents log transformed drug dose concentrations (μM). B) Half maximal inhibitory concentrations (IC50) for each of the drugs with or without radiotherapy were obtained using Prism (Graphpad, USA). A statistically significant difference in radiosensitisation across all three drugs (radiotherapy effect; Two-way ANOVA test*, p<0.0001) and between the different drug doses (drug effect: p<0.0001*). C) Results from western blot analysis of protein extracted from HCT116 following a 72 hour incubation with or without the three drugs (5FU-5μM; Apitolisib-100nM, Dactolisib-100nM), and with or without a single 5Gy dose of radiotherapy. Image densitometry was performed and relative pAKT expression has been demonstrated normalised to control (containing 0.1% DMSO). Protein was extracted two-hours after irradiation. Irradiation alone led to a significant increase in AKT phosphorylation at two-hours (p=0.027; two-tailed T-test**). Treatment with Dactolisib and Apitolisib with or without radiotherapy led to complete inhibition of AKT phosphorylation across three independent repeat experiments. API-Apitolisib; DACTO – Dactolisib; RT – radiotherapy; CI – confidence interval, pAKT –phosphorylated AKT, DMSO - Dimethyl sulfoxide.

Similarly, Cmax of 0.100μM, 0.220μM and 0.520μM were reported following the administration of 200mg, 400mg or 800mg of oral dactolisib to patients, respectively.^32^ Western blot analysis of phospho-AKT (p-AKT, Ser473) revealed that treatment with a single 5 Gy radiotherapy fraction led to a significant increase in AKT phosphorylation in HCT116 cells at two hours post-irradiation (*p*=0.027; Figure 6C). Standard treatment with 5μM of 5FU alone or 5μM 5FU with a single 5 Gy radiotherapy fraction did not significantly suppress AKT phosphorylation (*p*>0.05).

However, following treatment with 0.1μM of dual PI3K / mTOR inhibitors (dactolisib or apitolisib) there was complete suppression of AKT phosphorylation. AKT phosphorylation remained completely suppressed following administration of radiotherapy to HCT116 cells in the presence of dual PI3K / mTOR inhibitors.

### 2.5 Dual PI3K/mTOR inhibitors enhance radioresistant PDO response to radiation

We then investigated the radiosensitising potential of the dual PI3K/mTOR inhibitors on PDOs (Figure 7). When administered as single agent to radioresistant PDOs, both drugs where less efficient at reducing cancer cell viability compared to when in combination with irradiation (Figure 7A), providing a significant radiosensitising effect. The IC50 following apitolisib-only treatment for lines 557 and 653 were 5.0μM and 3.6μM respectively; combined with radiotherapy the IC50 was 1.3μM and 0.7μM. The IC50 following dactolisib-only treatment for lines 557 and 653 were 0.5μM and 0.3μM respectively; combined with radiotherapy the IC50 was 0.1μM and 0.04μM. (Figure 7B).

**Figure 7:**
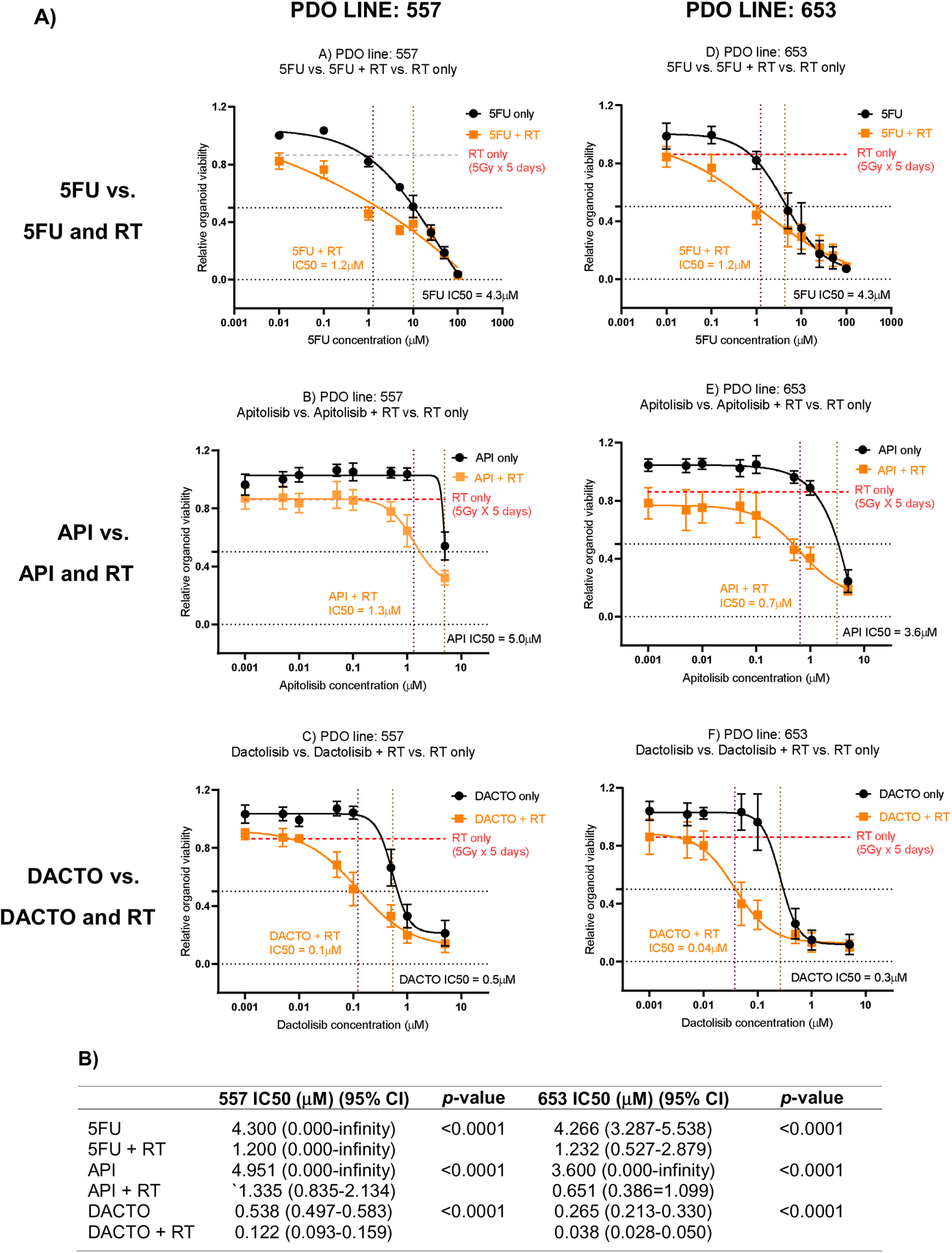
Radiotherapy resistant PDO lines 557 and 653 treated with 5FU, Apitolisib and Dactolisib with or without radiotherapy. A) PDO lines were plated at an estimated 5000 cells / well and treated with was treated with eight different concentrations of 5FU, Apitolisib (API) or Dactolisib (DACTO), with or without radiotherapy (5 Gy fractions over 5 days). End-point cell viability was assessed using CellTiter-Glo® 3D (Promega, USA) and chemiluminescence. The data is presented normalised to control (containing 0.1% DMSO). The x-axis represents log transformed drug dose concentrations (μM). B) Half maximal inhibitory concentrations (IC50) for each of the drugs with or without radiotherapy were obtained using Prism (Graphpad, USA). A statistically significant difference in radiosensitisation across all three drugs (radiotherapy effect; Two-way ANOVA test*, p<0.0001) and between the different drug doses (drug effect: p<0.0001*).

## 3. Discussion

LARC patients who achieve pCR following CRT demonstrate improved disease-free and overall survival compared to non-responders or partial responders.^7, 33^ Furthermore, pre-operative detection of a complete clinical response (cCR) following CRT, enables the use of “watch and wait” strategies in the management of LARC patients^34^, potentially avoiding the morbidity of major surgery.^35^ The role of CRT in the treatment of early RC is also being investigated.^36^ Hence, the need to unravel the biological mechanisms underlying radioresistance in LARC in order to improve cCR or pCR rates. To address this, we used PDOs given their proven ability to function as tumour avatars that recapitulate the parent tumour genome and mimic clinical response to therapy.^27, 28^ After mapping the PDO mutation profiles (Figure 1), their radiation response was assessed (Figure 2). Four lines were deemed radiosensitive (411, 064, 389, 884) and two radioresistant (557, 653).

We found *KRAS* mutations (*G12D* and *G13R*) in the radioresistant PDOs, and analysis of their transcriptome showed enrichment of KRAS downstream target genes (Figure 3C). *KRAS* mutation status is a predictor of poor response to anti-EGFR antibody therapy in metastatic CRC, and its correlation with RC response to CRT has been widely studied although a consensus was not reached.^37–39, 40–43^ A systematic review and meta-analysis of 696 patients concluded that *KRAS* status was not predictive of tumour downstaging following CRT in LARC patients.^44^ However we hypothesise that significant molecular heterogeneity exists within these studied tumours, and that the presence of a KRAS mutation at a level undetectable by conventional sequencing, may lead to the domination of this KRAS mutation clone when irradiated.

After evaluating PDOs response to radiation, we aimed to identify novel genes and pathways that promote radiation resistance by comparing the pre-treatment transcriptomes of radiation-sensitive and -resistant PDOs. The most significantly upregulated genes found in radioresistant PDOs (Figure 3B) have known functions of modulating cell response to radiation or are associated with cancer progression.

The genes *CAV1* and *SCARA3* have been previously described to be involved in ionizing radiation response or resistance. *CAV1* encodes an integral membrane protein that has been found over-expressed or mutated in solid human tumours.^45^ In pancreatic and lymphoblastoid cancer cells, CAV1 acted as a pro-survival factor mediating resistance to the cytotoxic action of ionizing radiation.^46^ Additionally, CAV1 overexpression was found in triple negative breast cancer cell lines and its expression levels correlated with the sensitivity to ionizing radiation.^47^ Scavenger receptor class A member 3 (SCARA3) protects against oxidative stress induced cell killing and can serve as predictor of progression and therapeutic response in multiple myeloma.^48^ Other significantly over-expressed genes in radioresistant PDOs – namely *PLK2*, *UBASH3B*, and *NTN1* – have been associated with cancer progression. Ou et al. found that colorectal tumours expressing high levels of Polo-like kinase 2 *(PLK2)* display lower disease-free survival.^49^ The authors further showed *PLK2* promotes tumour growth and inhibits apoptosis of CRC cells *in vitro* and *in vivo* by binding to *FBXW7* and subsequently promoting its degradation, which in turn leads to the stabilization of Cyclin E. *UBASH3B* gene encodes a ubiquitin receptor protein that controls spindle assembly checkpoint silencing and faithful chromosome segregation. Increased cellular levels of this gene trigger premature and uncontrolled chromosome segregation.^50^ High expression of netrin1 (NTN1) – a laminin-related secreted protein – is found in at the base of colon crypts, where the intestinal stem cells are located.^51^ NTN1 triggers anti-apoptotic signalling on cells presenting the dependence receptors DCC and UNC5H.^52^ Several authors have attributed NTN1 pro-tumorigenic effect to activation of PI3K/AKT pathway.^53–55^

Principal component analysis of pre-treatment PDO transcriptomes showed general sample diversity reflecting patient inter-tumour variability and genetic heterogeneity as seen in our single cell transcriptomic data. Notwithstanding, it grouped the PDOs in two clusters according to radiosensitivity (Figure 3A).

Radioresistant PDOs displayed upregulation of the epithelial-mesenchymal transition (EMT) pathway and the PI3K/AKT/mTOR pathway (Figure 3E), both of which have been associated with radioresistance in RC.^56–58^ The role of EMT is well established in cancer metastasis and therapy resistance.^59^ This could suggest rectal tumours enriched with cancer cells in a mesenchymal state are more radioresistant. Furthermore, a close association between PI3K/AKT/mTOR pathway activation and EMT pathway protein expression leading to radioresistance has been previously described.^60, 61^ Furthermore, the results from PDO single cell sequencing revealed upregulated PI3K/AKT/mTOR pathway and DNA damage repair genes following irradiation (Figure 4E). Genome wide CRISPR-Cas9 knockout screen demonstrated that this pathway was crucial for the survival of irradiated CRC cells (Figure 5C).

CRT-resistant tumours demonstrate the ability to evade apoptosis, enhanced DNA double-strand breaks repair, changes to cellular metabolism, resistance to hypoxia, poor inflammation and abundance of tumour stem cell populations.^62^ The PI3K/AKT/mTOR pathway regulates several of these functions.^63^ Pathway activation leads to several downstream effectors responsible for cell proliferation, migration, growth and DNA repair.^64^ The pathway is implicated in the pathogenesis as well as CRT resistance in various cancers.^65, 66^ Mutations within genes coding for components within this pathway are frequently detected in malignancy.^67^

Several pharmacological inhibitors targeting the PI3K/AKT/mTOR pathway are clinically available and have been tested for cancer treatment,^68, 69^ and the radiosensitising potential of these inhibitors is currently being assessed in several tumour types.^70^ In CRC, studies using cell lines and cell line-derived xenografts have shown that PI3K/mTOR inhibitors dactolisib and PI-103 improve the efficacy of ionizing radiation.^71–74^ Long-term administration of PI-103 before irradiation may cause reactivation of PI3K and MAPK pro-survival pathways,^74^ we opted to exclude it from our assays. Preliminary experiments showed no increase in organoid radiation sensitivity when co-treated with Sirolimus or everolimus (data not shown).

Furthermore, early clinical trials evaluating the radiosensitising effects of selective mTOR inhibitors such as everolimus, rapamycin or sirolimus failed to increase pCR rates.^75–77^ Emerging preclinical evidence points towards dual pathway component inhibitors as effective radiosensitisors,^70^ guiding our option of testing dual PI3K and mTOR inhibitors (apitolisib and dactolisib). In addition, dactolisib has safely undergone phase Ib clinical trials in humans in various cancers.^31, 32, 75^ Apitolisib has completed phase II clinical trials in human cancer patients.^76, 77^

Isogenic *KRAS* HCT116 CRC cell line was chosen for our experiments given it was in stable culture without requiring passage during 5 days of irradiation and has been widely used in CRC *in vitro* research,^78^ including those evaluating response to radiation.^71, 74^ In our assays, dactolisib showed radiosensitising effects in HCT116 and radioresistant PDOs, with an IC50 lower than the previously published Cmax in humans. Conversely, the IC50 for apitolisib-treated radioresistant PDOs exceeded the drugs Cmax in humans, rendering not useful for clinical practice. The different pharmacological properties of the two drugs may explain the differences observed between apitolisib and dactolisib response in PDOs. Our results also demonstrated AKT phosphorylation when HCT116 was treated with clinically comparative doses of radiotherapy (Figure 6C). This was in keeping with findings published by Chen et al.^71^ We observed complete p-AKT suppression in apitolisib and dactolisib-treated cells (Figure 6). Suppressed AKT phosphorylation could lead to inhibition of several downstream pathway functions such as DNA repair, cell survival, proliferation and migration,^70^ in turn would lead to radiosensitisation.

We acknowledge the limited availability of patient samples and model derivation success (approximately 40%) constituted a limitation to this study resulting in a reduced sample size. Nevertheless, the biological pathways upregulated in radioresistant PDOs identified in this study were in keeping with the literature.

Subsequent studies encompassing larger and more homogeneous cohorts are warranted to further evaluate the effects of radiotherapy and pathway blocking drugs on PI3K/AKT/mTOR pathway downstream effectors.

In summary, this study utilised state-of-the-art patient-derived primary cell culture models to investigate the biological mechanisms driving radiation resistance in RC. This led to the identification of several genes which could serve as targets for novel therapies and biomarkers for prognostication. Furthermore, this study showed that biological pathways such as EMT and PI3K/AKT/mTOR are upregulated in radioresistant tumour cells. We demonstrated that dual PI3K/mTOR inhibitors improve radiotherapy efficiency on radioresistant PDOs, warranting further studies to validate the use of these drugs in the clinical setting.

## 4. Methods

### 4.1 Patient samples

Patients undergoing surgery for primary colonic or rectal adenocarcinoma were recruited from Queen Elizabeth Hospital Birmingham, UK. Fresh tumour tissue specimens sampled from surgical resection by pathologists were used for tumour organoid derivation. A total of 6 PDOs successfully passed the derivation and propagation stage (Table 1): three from distal adenocarcinomas originating within the rectum (884, 653) or rectosigmoid junction (064), two from the transverse colon (389, 411) and one from the ascending colon (557). Four were treatment naïve (064, 389, 411, 557), whereas 884 received SCRT and 653 received CRT. Anonymised clinicopathological data were obtained from the local Human Biomaterials Resource Centre in Birmingham.

**Table 1:** Clinicopathological characteristics of the parent tumours from which the six PDO lines were derived

| ID | Origin | Age | M/F | Diagnosis | Nx | TRS | TNM | MMR | KRAS | BRAF |
| --- | --- | --- | --- | --- | --- | --- | --- | --- | --- | --- |
| 884 | Rectum | 75 | M | Adenocarcinoma | SCRT | TRS-3 | pT3, N2a, M0 | No loss | wt (N-RAS-mut) | wt |
| 064 | Rectosigmoid | 79 | M | Adenocarcinoma | NA | NA | pT2, N1, M0 | No loss | wt | wt |
| 389 | Colon | 82 | M | Adenocarcinoma | NA | NA | pT4a, N2a, M0 | No loss | wt | wt |
| 411 | Colon | 74 | F | Adenocarcinoma | NA | NA | pT3, N2b, M0 | No loss | wt | mut |
| 653 | Rectum | 55 | F | Adenocarcinoma | NCRT | TRS-2 | pT2, N0, M0 | No loss | mut | wt |
| 557 | Colon | 76 | F | Adenocarcinoma | NA | NA | pT3, N0, M0 | Loss of MLH1, PMS2 and MHS6 | mut | Not available |

### 4.2 Organoid and cell culture

Tumour resection specimens were processed into single-cell suspension by mechanical and enzymatic digestion, mixed with Matrigel® (Corning, USA) and incubated in 50µl droplets on 24-well plates, as previously described by Sato et al.^79^ Organoid culture media (OCM) Human Intesticult™ (Stemcell Technologies Inc, Canada) containing 0.2% Primocin™ (Invivogen, USA) and passaged once confluent. Isogenic *KRAS* HCT116 (Horizon Discovery, Cambridge, UK) and C80 (#12022904-1VL, Sigma Aldrich, USA) CRC cell lines were cultured in McCoy’s 5a Medium (Gibco, USA) with 10% fetal bovine serum (FBS, Gibco, USA), Iscove’s modified Dulbecco’s media (Gibco. USA) containing 10% FBS, 2mM L-Glutamine (Gibco, USA) and Eagles modified essential medium (Gibco, USA) with 2mM L-Glutamine containing 1% non-essential amino acids (Sigma-Aldrich, USA), respectively, and one hundred units/ml penicillin and 100 μg/ml streptomycin (Gibco, USA). Cells were passaged when sub-confluent at a split ratio of 1 in 3. All cultures were tested for mycoplasma using EZ-PCR Mycoplasma Test Kit (Biological Industries, Israel).

### 4.3 Organoid irradiation experiments

Suspensions of 5,000 cells from each PDO line were added to Matrigel-coated wells of 96-well plates. Different doses of radiation (2, 5, 10, 20 and 40 Gy) were delivered over five consecutive days using a CellRad irradiator (Faxitron Bioptics, AZ, USA). A minimum of 4 replicates were used per experimental condition. PDOs were incubated for additional 5 days to allow cell recovery to assess cell viability within the range of sensitivity of the cell-viability protocol. Cell viability was assessed by luminescence following the CellTiter-Glo® 3D Cell Viability Assay protocol (Promega, USA). Data were normalized to the PDO wells treated with 0 Gy as 100% viability to calculate relative cell viability of the irradiated wells.

### 4.4 Immunohistochemistry

PDOs were released from Matrigel® drops, fixed in 4% paraformaldehyde and cast in 2% agarose pellets. These were then embedded in formalin-fixed paraffin embedded (FFPE) blocks using standard protocols. Slides with 5µm organoid sections were stained for hematoxylin and eosin (Sigma Aldrich, USA) for morphology assessment. FFPE sections of all PDOs and their respective tumour of origin were stained with antibodies targeting pan-Cytokeratin (ab27988, Abcam, UK) and CDX2 (9272S, Abcam, UK) following standard protocols. The primary antibody dilution was 1/500 and 1/100, respectively, and both were incubated for 1 hour.

### 4.5 Targeted DNA sequencing

DNA from PDO cells was extracted using the AllPrep DNA/RNA Mini Kit (Qiagen, Germany). Library preparation was performed using a QIASeq™ custom targeted 30 gene DNA panel (Qiagen, Germany), detailed in Supplementary Table 1. Sequencing was performed using a MiSeq v2 (Illumina, USA), 300 cycle flow cell paired-end at 500X coverage on a MiSeq™ (Illumina, USA) next generation sequencing (NGS) platform.

### 4.5 RNA sequencing

RNA from PDO cells was extracted using the AllPrep DNA/RNA Mini Kit. Ribosomal RNA (rRNA) was depleted using the rRNA depletion Kit (Human/Mouse/Rat) (New England BioLabs [NEB, USA]). Library preparation for total RNA sequencing was performed using NEBNext^®^ Ultra^TM^ II RNA Library Preparation Kit for Illumina (NEB, USA). Sequencing was performed using a NextSeq HIGH 150 (Illumina, USA) flow cell on the NextSeq™ 500 (Illumina, USA) NGS platform.

### 4.6 Single cell RNA sequencing

Irradiated (5 Gy/day over 5 days) and non-irradiated PDOs were dissociated into single cells after a 6 day recovery period. Libraries were prepared aiming for 7000 cell recovery, using 11 cycles of amplification of cDNA and 12 cycles for indexing PCR using the Chromium single cell 3’ kit (10X Genomics, USA). Sequencing was performed using a NextSeq™ 550 HIGH 150 flow cell and NGS platform.

### 4.7 Genomewide CRISPR-Cas9 knockout

The GeCKO pooled library (#1000000048, Addgene, USA) was amplified as per the manufacturer’s instructions. Sufficient single guide RNA (sgRNA) coverage of the resulting library was confirmed on the Illumina MiSeq™. Lentivirus generation and subsequent transduction and selection of C80 cell line was performed as described by Joung et al.^80^ The CRC cell line used in the CRISPR-Cas9 experiments, C80 contained *APC*, *KRAS*, *SMAD4* and *TP53* mutations. Two cell populations were maintained, each with a coverage of 500 for each sgRNA (6.1X10^7^ cells). One population was treated with 5 Gy/day radiotherapy over 5 days and the second population remained untreated. Allowing for a 14 day, post-treatment period for cell recovery, 6.1×10^7 cells from each group were harvested, RNA extracted and library preparation performed with individual barcoding of each group. NGS was performed using Illumina NextSeq™.

### 4.7 In vitro cell line and PDO chemoradiotherapy assays

HCT116 (1000 cells/well) and two radioresistant PDO lines (organoids equivalent to 5000 cells/well) were treated with 5-fluorouracil (5FU, PanReac-AppliChem, USA) or dual PI3K / mTOR inhibitors apitolisib (GDC0980, Adooq Bioscience, USA) or dactolisib (BEZ235, Selleckchem, USA), in 0.1% DMSO, with or without radiotherapy (5 Gy over 5 days for PDO lines and 1 Gy over 5 days for HCT 116). Ninety six well assay plates were prepared. Drug treatment commenced on the third day after plating and radiotherapy on the fourth day. During the experiment, wells were replenished with 100μl of culture media (with or without drug) every 72 hours. The treatment ended on ninth day when all wells were replenished with 200μl of fresh culture media and incubated for a further 5 days to allow cell death or recovery. End-point viability was assessed with chemiluminescence using CellTiter-Glo 2.0 Assay (Promega, USA) and CellTiter-Glo® 3D Cell Viability Assay for cell and PDO lines, respectively.

### 4.8 Western blots

The protein extracted from HCT116 cells following drug or radiotherapy treatment, was loaded at 20μg on to precast gels. Western blots were developed using protocols from Abcam Plc (UK) and primary antibodies Phospho-AKT Ser473 (4060S, Cell Signalling Technology, USA), AKT (9272S, Cell Signalling Technology, USA) and GAPDH D16H11 (5174S, Cell Signalling Technology, USA).

### 4.9 Data analysis

Prism V8 (Graphpad, USA) was used to generate dose-response graphs. Half maximal inhibitory concentrations (IC50) with 95% confidence interval (CI) were determined using linear regression. Two-way ANOVA mixed model analyses were performed using Prism V8 to assess significance in efficacy of radiotherapy or drug treatment on cell line and PDOs. Western blot experiments were quantified using ImageJ (https://imagej.nih.gov/ij/). A student t-test was performed using SPSS (IBM, USA) to assess significance of AKT phosphorylation or suppression. QIASeq data were analysed using the Biomedical Genomics Workbench (Qiagen, Germany). Mutation profile heatmap was generated using Oncoprint Chart. RNA NGS data were used to identify differentially expressed genes using pre-designed bioinformatic pipelines on Flow® (Partek, UK). DESeq2 was applied to evaluate the differential gene expression during the transition of the two developmental stages using DEBrowser v1.16.1.^81 82^ Genes were considered differentially expressed when they had an adjusted p-value ≤ 0.01 (FDR); fold-change ≤ −2. Differentially expressed genes were plotted using the EnhancedVolcano.^83^ Over-representation analysis using gene sets from the MSigDB v7.4 (https://www.gsea-msigdb.org/gsea/index.jsp) was performed against differentially expressed genes using the R package clusterProfiler.^84^ The R packages were run on R Studio Desktop V1.3 (R Studio, USA) with R V4.0.2 (r-project.org, USA).

For single cell sequencing, read matrices were imported into R 4.0.4 (Lost Library Book) and BioConductor and the GRCh38 reference genome. Cells were filtered such that cells with > 5% mitochondrial reads or had < 200 features were excluded. Data were log normalised and non-linear dimensionality reduction performed via UMAP, and differential expression markers between clusters were found using the *FindMarkers* function of Seurat. Cell identity based on differential expression clusters was performed by a combination of KEGG pathway analysis and manual annotation based on the literature. Cell populations were segregated into those from irradiated organoids and those from control organoids and Scalable Bayesian Boolean Tensor Factorisation (SBBTF) using Python 3.7.3, which binarised gene expression with outcome.

Following the CRISPR/CAS9 screen, sequenced reads were analysed using the MaGECK pipeline. Briefly, sgRNA counts for each cell line were performed using the mageck *count* command and differential changes in sgRNA between cell types was carried out using the mageck *mle* command, using CNV data from the Depmap project to correct the sgRNA calls for alterations in CNV. Differential sgRNA data were processed using MaGECKFlute. Gene Ontology over-representation analysis was performed using the R package clusterProfiler. The ‘org.Hs.eg.db’ v3.13.052 genome-wide annotation was supplied for mapping Ensembl IDs to gene symbols.

Significantly over-represented KEGG and gene oncogenic terms were defined from molecular signatures database v7.4 using differentially expressed genes filtered by *p* < 0.05. The ‘dotplot()’ function was used to create visualisations of the top over-represented gene sets. Network modules of highly correlated genes was performed using the R package ‘WGCNA’ v1.70-353. The soft thresholding power was selected using the ‘pickSoftThreshold()’ function and scale-free topology criterion. An adjacency matrix was created, containing the Pearson’s correlation coefficients of every possible pair of genes. The adjacency matrix was used to create a topological overlap matrix (TOM). The corresponding dissimilarity matrix (1 - TOM), was used for gene hierarchical clustering and subsequent module detection using the Dynamic Tree Cut algorithm. Each module detected was named using a colour.

## Acknowledgements

We acknowledge the consultant body of the colorectal surgery department at the University Hospitals Birmingham NHS Foundation Trust (UHBNHSFT) for their support with patient recruitment. We are also grateful for the support of the team at the histopathology department and operating theatres at UHBNHSFT for their help with tissue sampling and specimen retrieval. We would like to extent our gratitude to the staff at The Human Biomaterials Resource Centre (BioBank), Birmingham for their help in sourcing the fresh tissue samples and anonymised clinical data for this research. We express our gratitude to Cancer Research UK for funding this research.

## Contributors

KW and JBS designed the experiments under the supervision of TI and ADB. KW, OJP, LT, AS, TS and RH conducted the wet lab experiments and immunohistochemistry. LT conducted the CRISPR-Cas9 experiments. KW, OJP, AS, LT, CB, VP, JS, CW, RT and TS were involved in the genomic work packages. KW, JBS, ME, CY and ADB were responsible for data analysis including bioinformatics and data interpretation. KW and JBS wrote the manuscript with the support of ME, LT, JS and ADB. The remaining authors were also involved in the editing and final review of the manuscript.

## Patient consent

Obtained

## Ethics approval

Project approval code 17-287, Human Biomaterials Resource Centre, Birmingham, United Kingdom; under ethical approval from North West - Haydock Research Ethics Committee (Reference: 15/NW/0079).

## Availability of data and materials

All relevant data and results included in this article have been published along with the article and its supplementary information files. Other relevant data can be obtained on reasonable request from the corresponding author.

---

**Supplementary Table 1:**
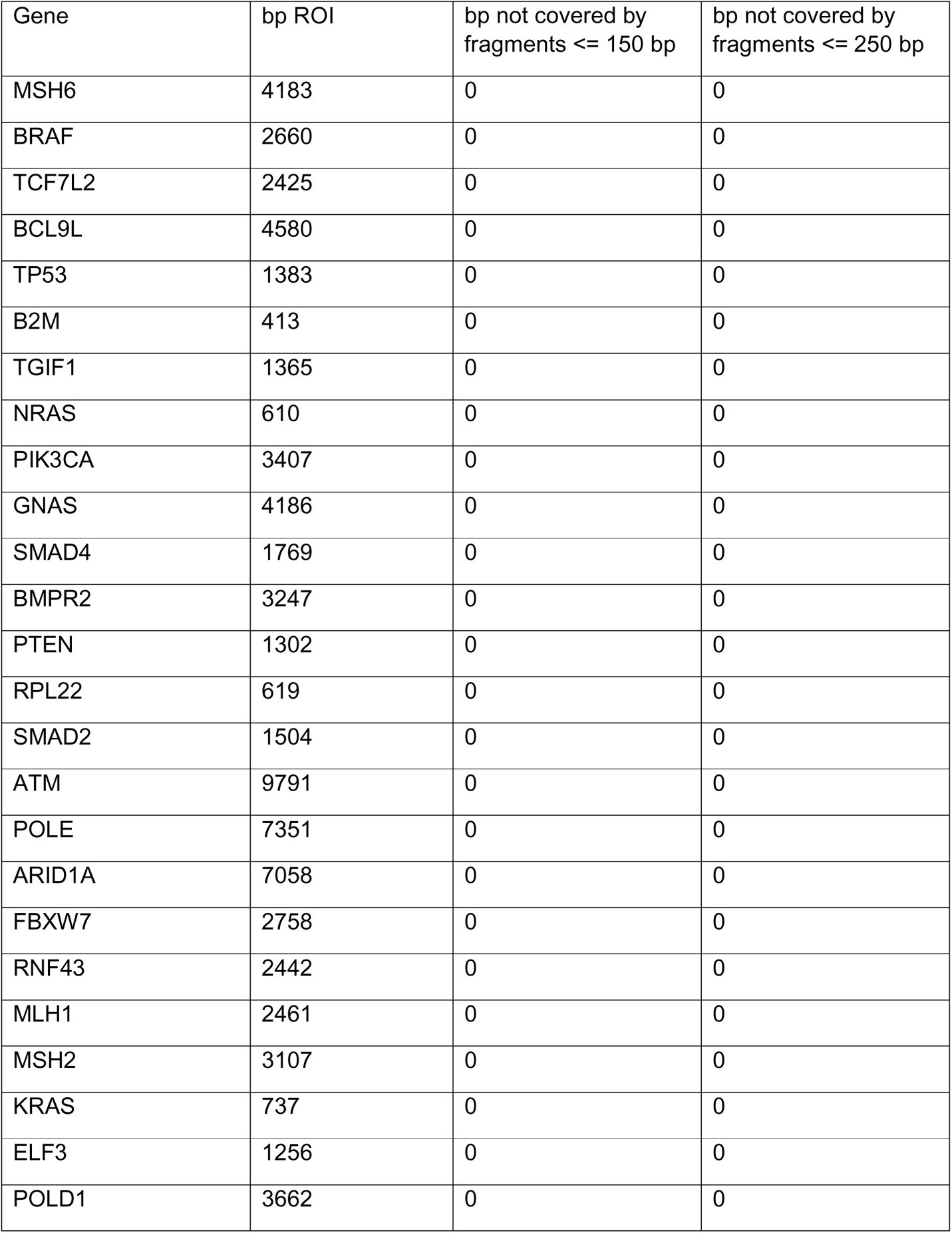
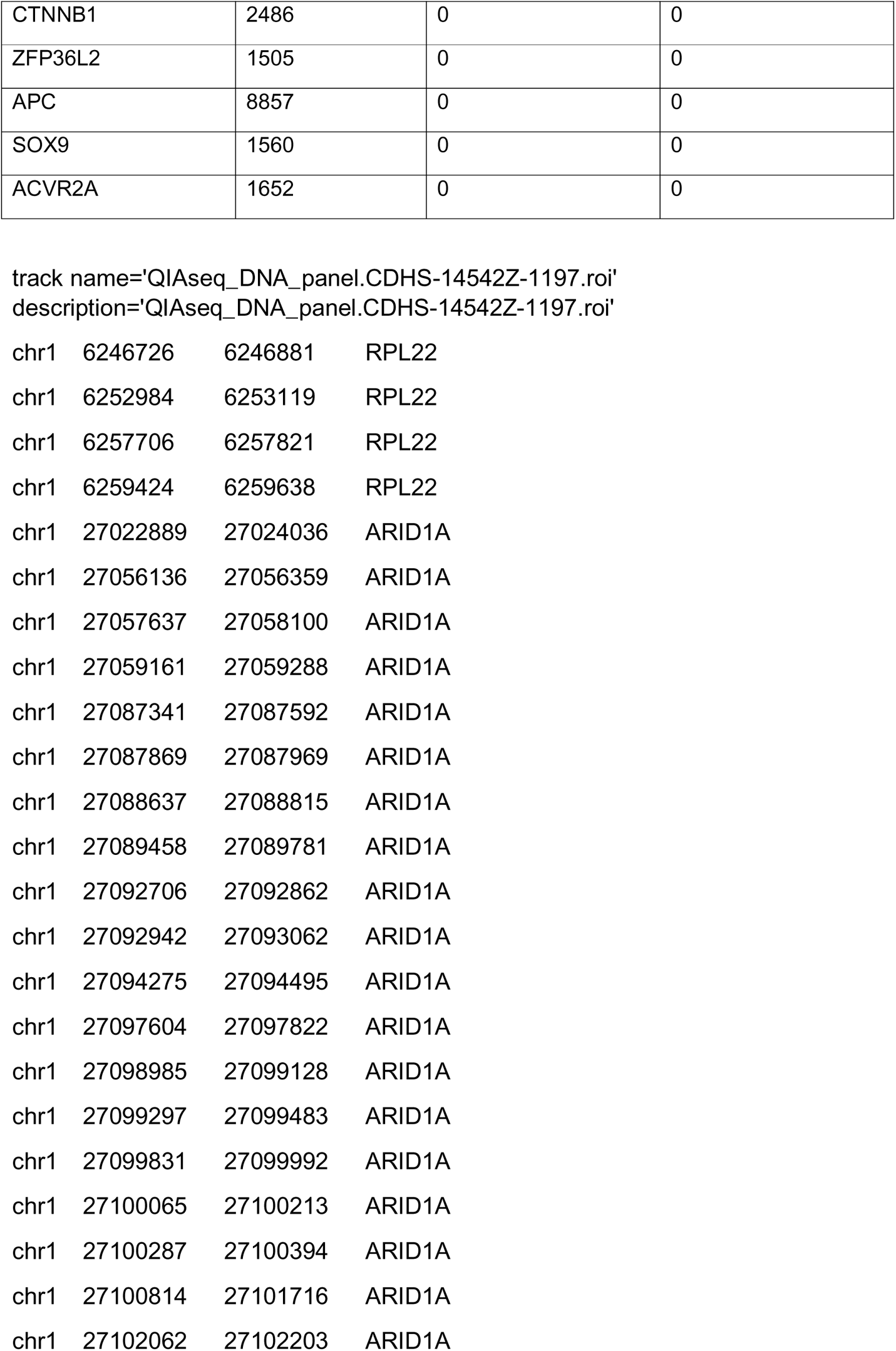
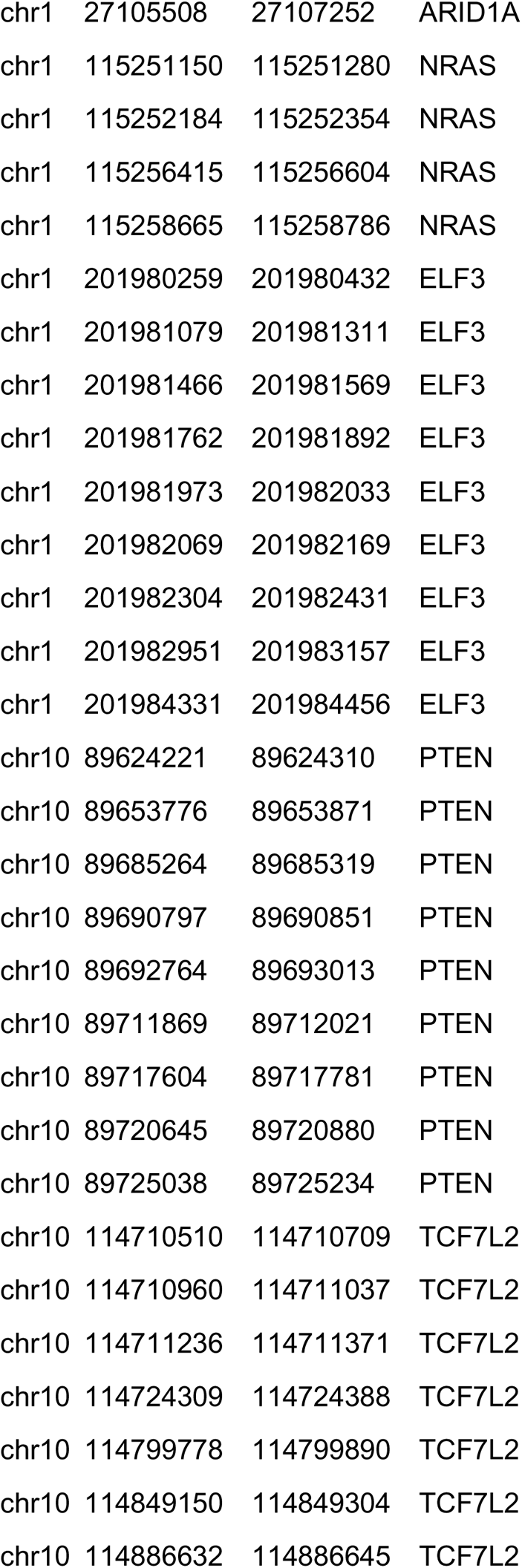
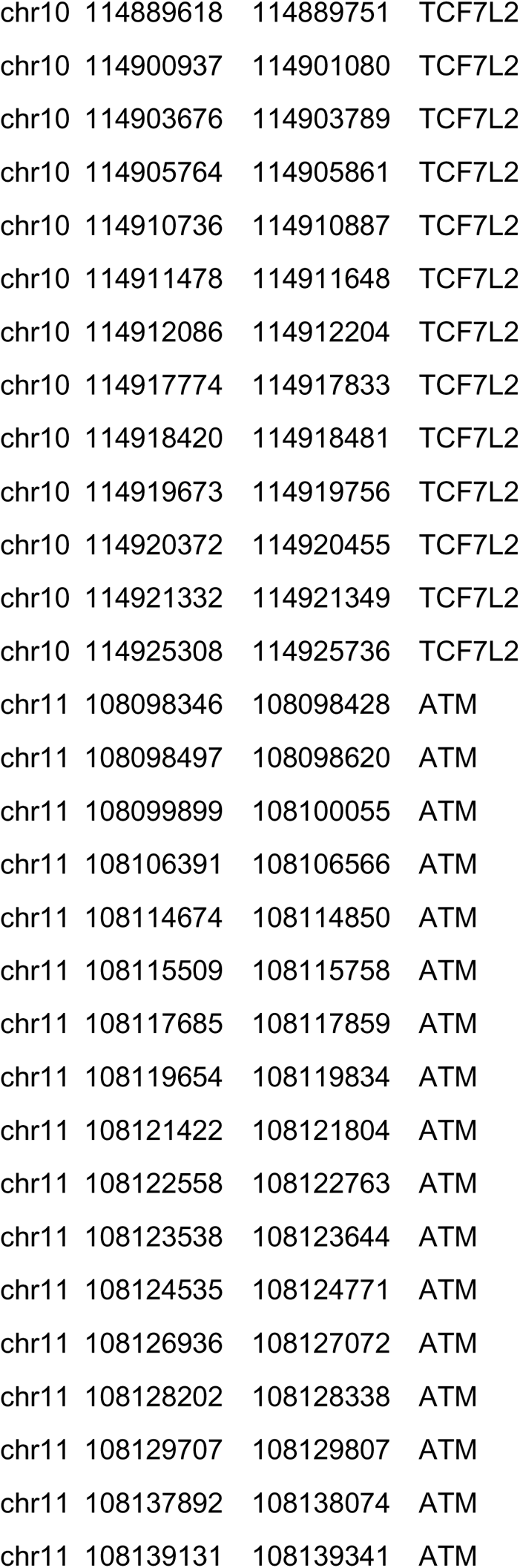
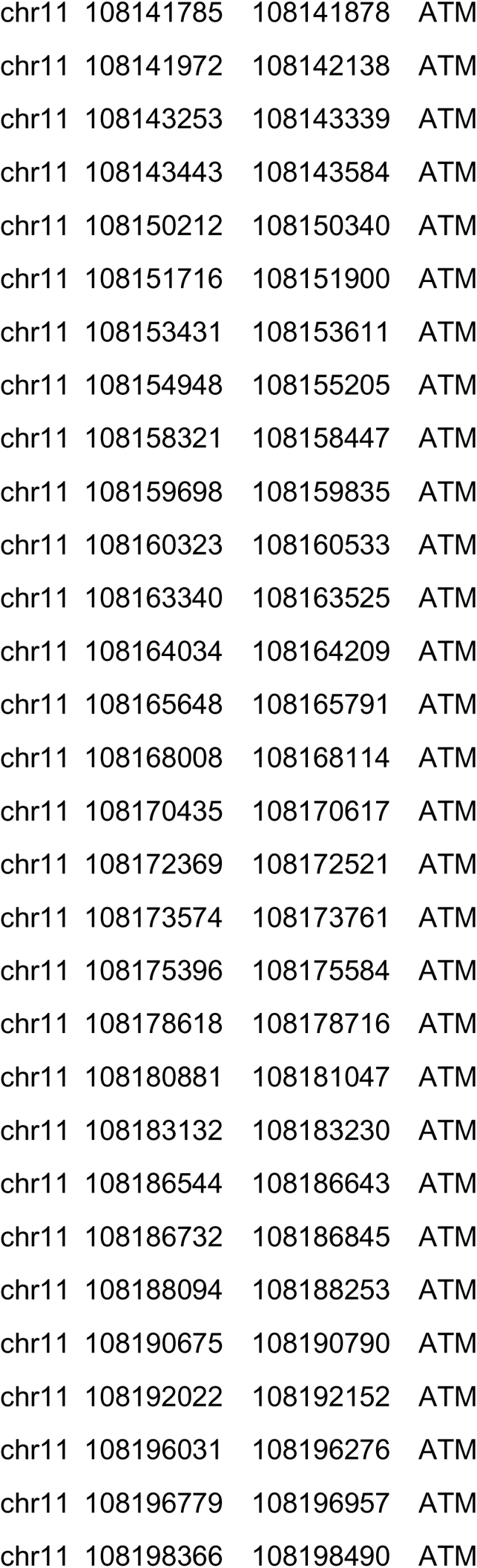
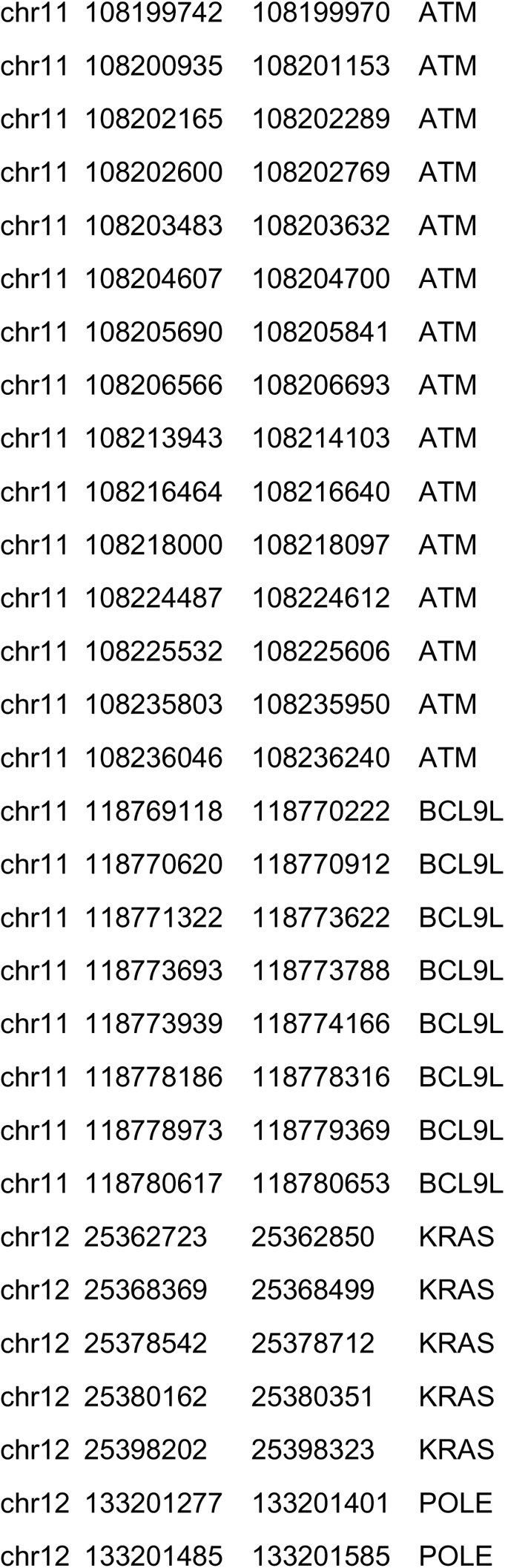
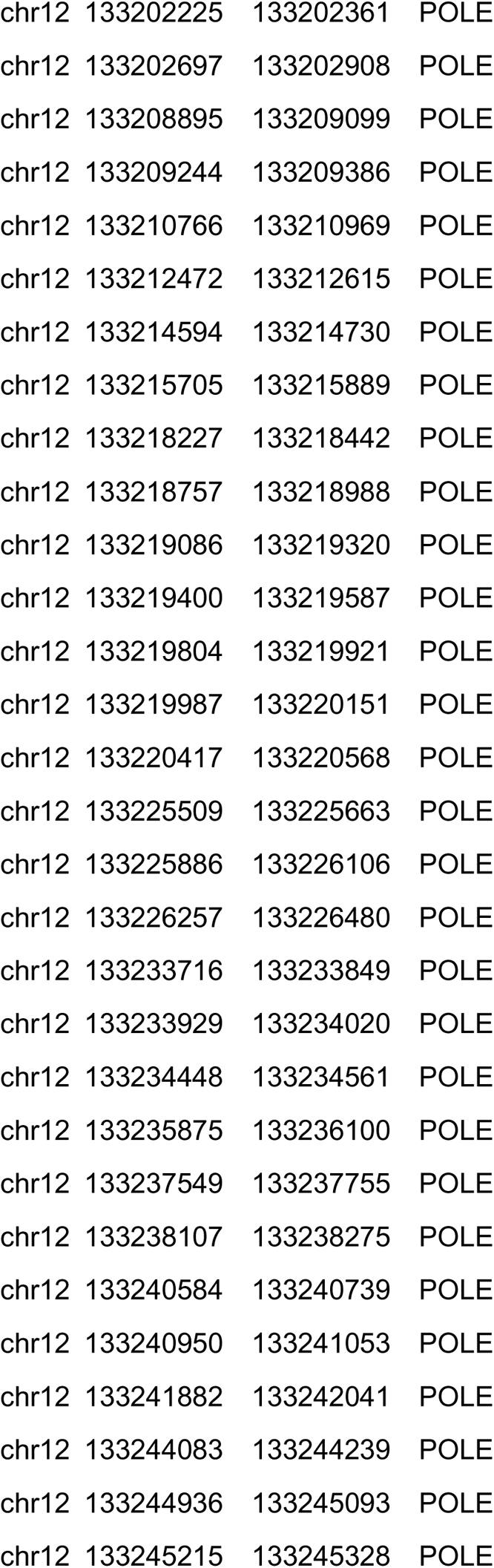
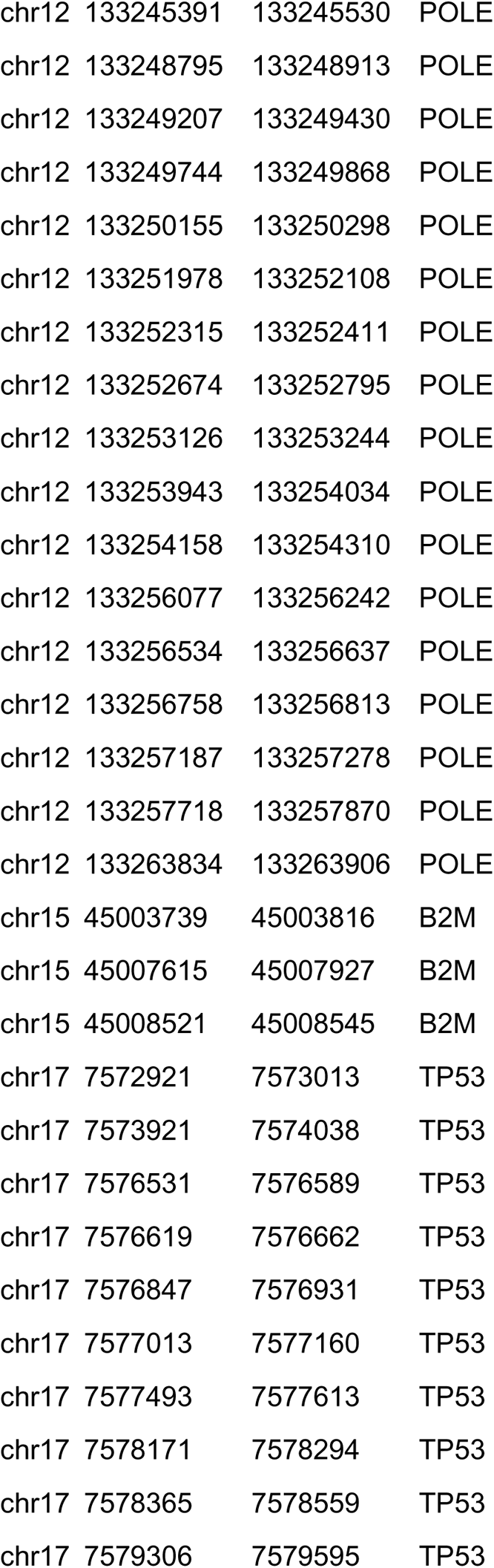
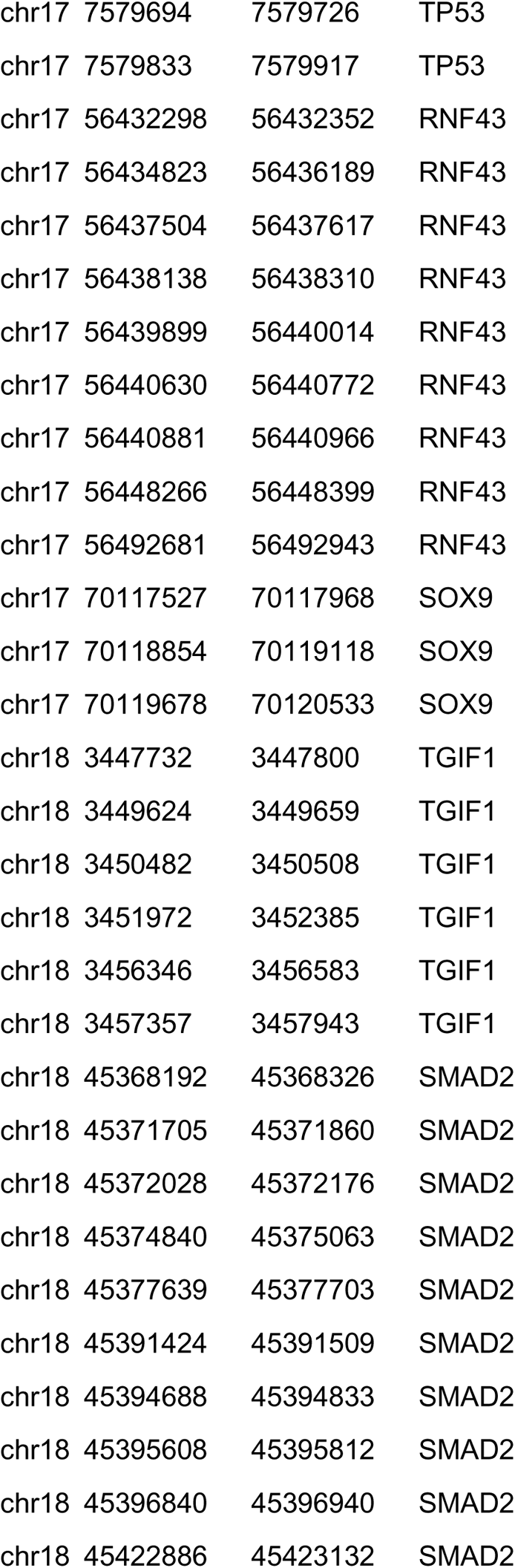
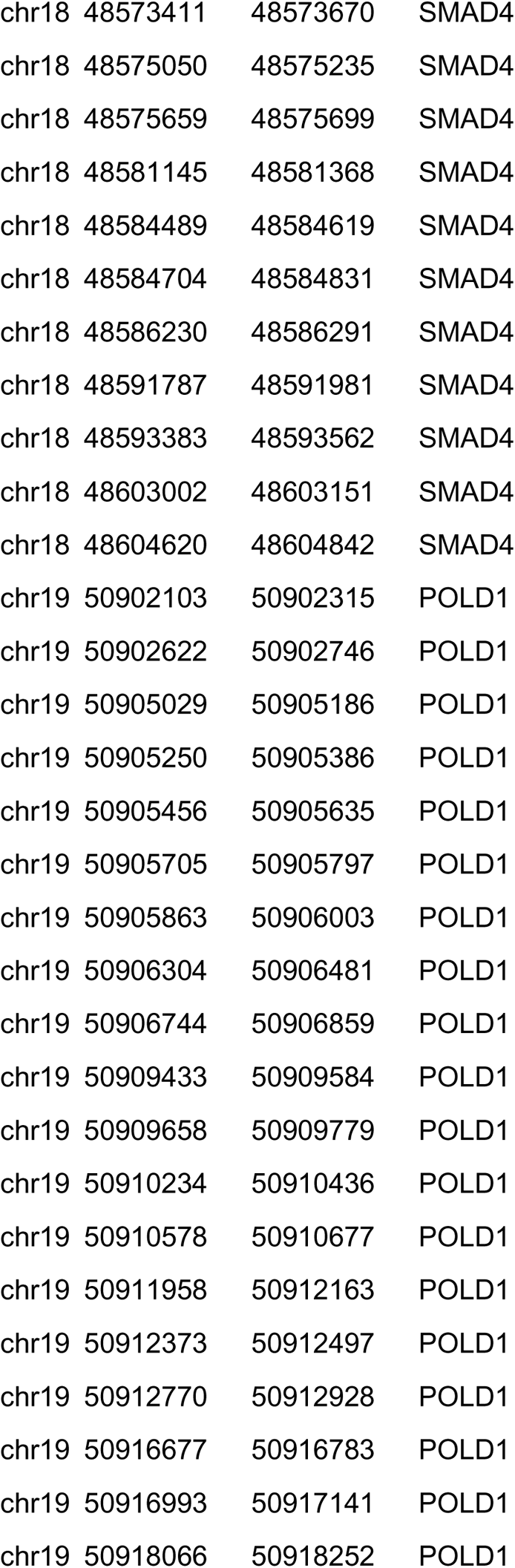
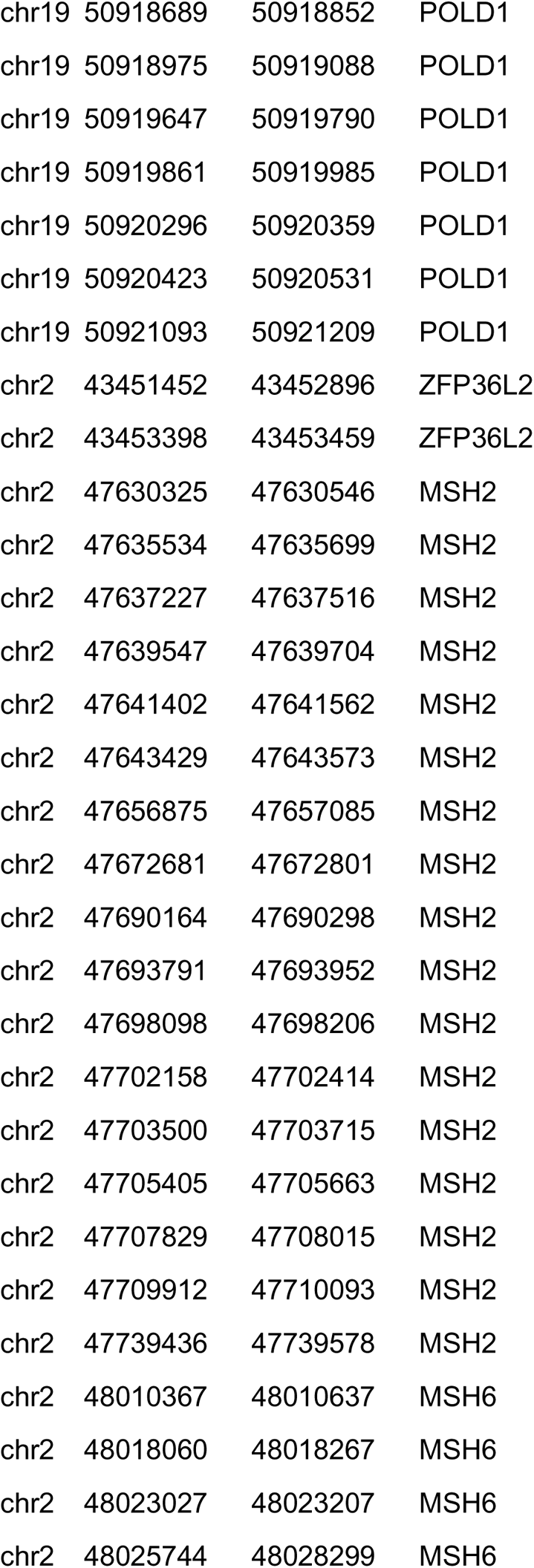
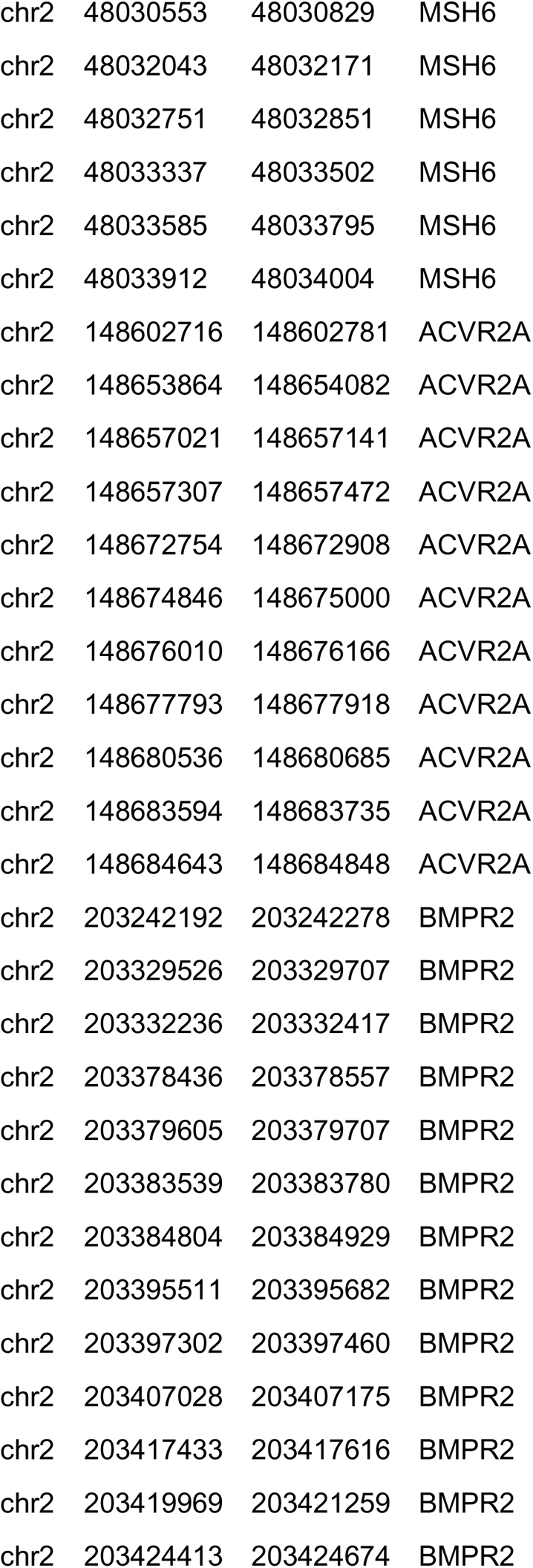
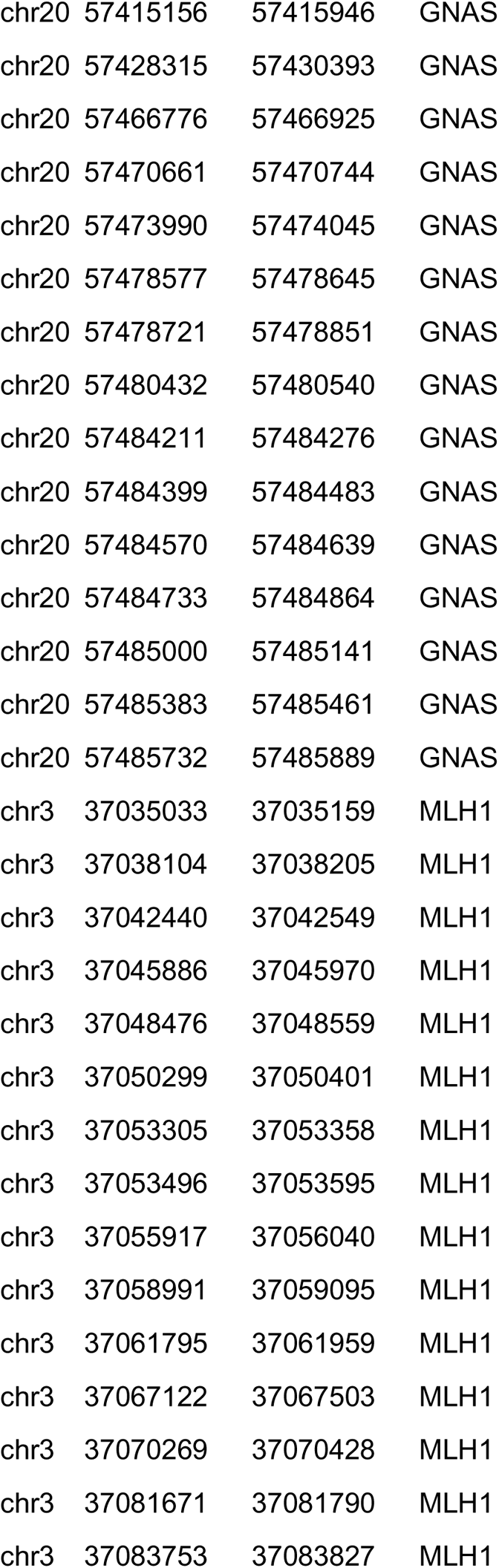
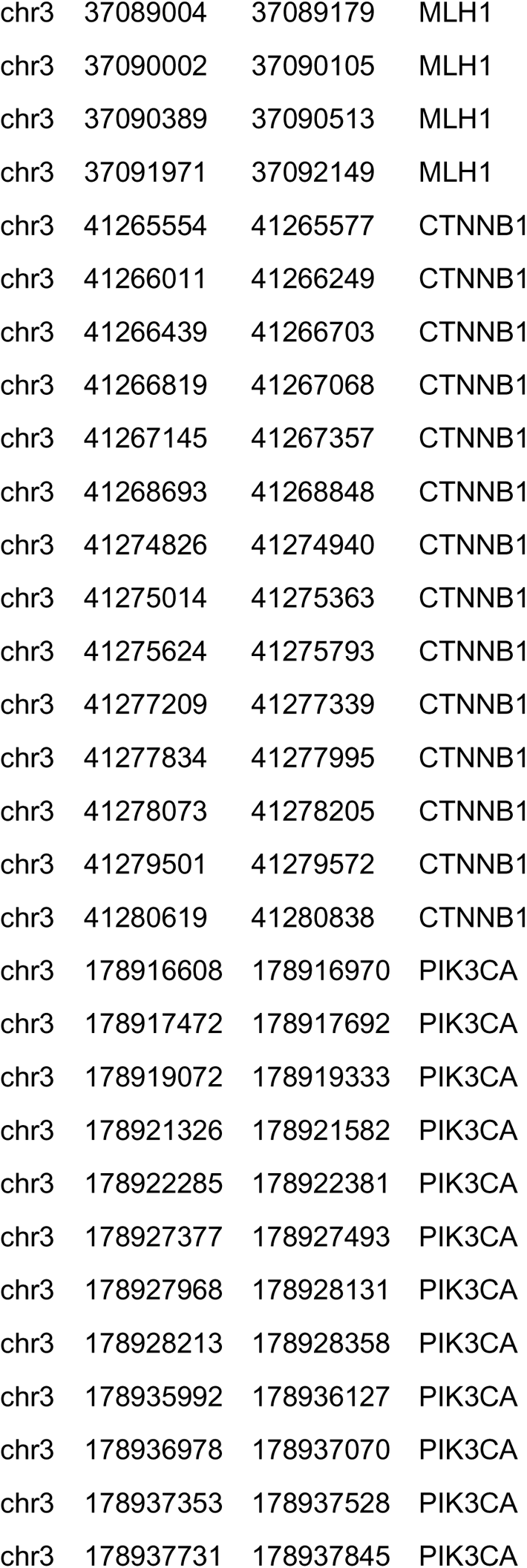
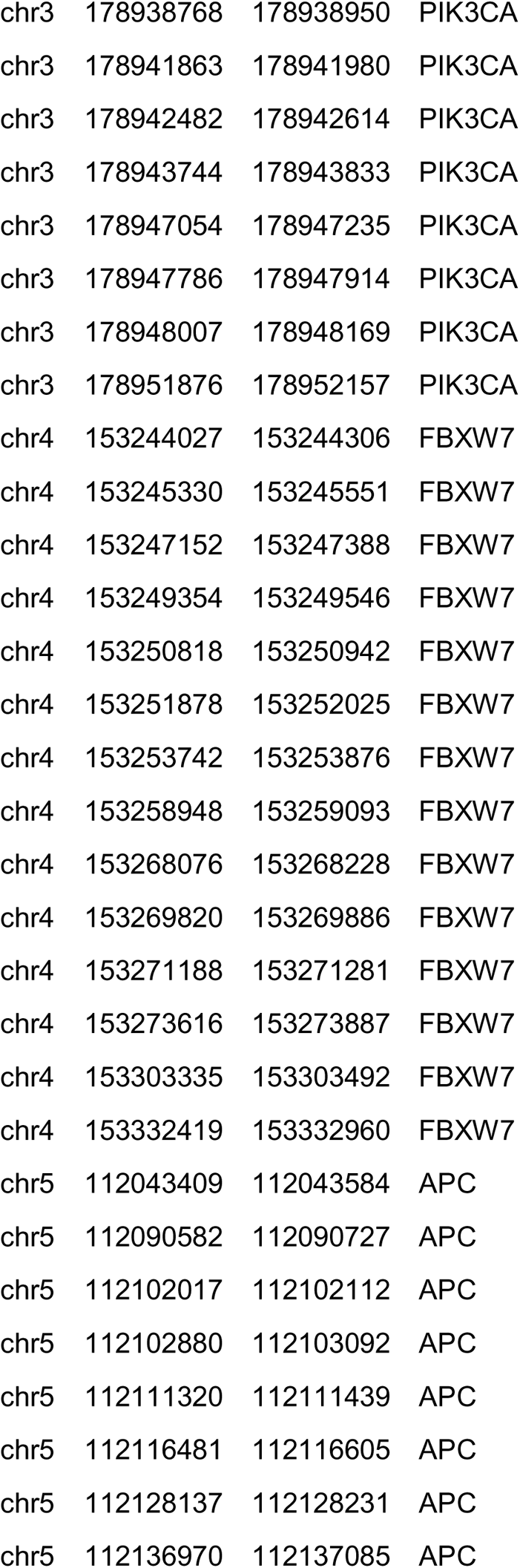
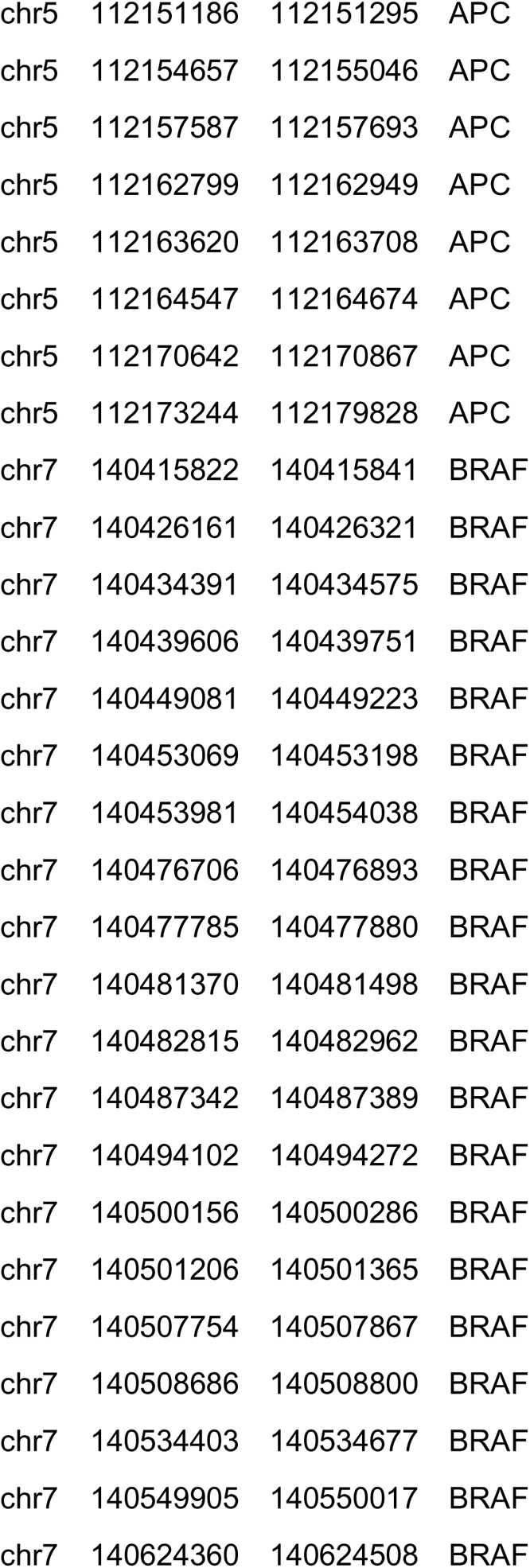
Genes tested using the QIASeq™ (Qiagen, Germany) custom targeted DNA sequencing panel

| Gene | bp ROI | bp not covered by<br>fragments <= 150 bp | bp not covered by<br>fragments <= 250 bp |
| --- | --- | --- | --- |
| MSH6 | 4183 | 0 | 0 |
| BRAF | 2660 | 0 | 0 |
| TCF7L2 | 2425 | 0 | 0 |
| BCL9L | 4580 | 0 | 0 |
| TP53 | 1383 | 0 | 0 |
| B2M | 413 | 0 | 0 |
| TGIF1 | 1365 | 0 | 0 |
| NRAS | 610 | 0 | 0 |
| PIK3CA | 3407 | 0 | 0 |
| GNAS | 4186 | 0 | 0 |
| SMAD4 | 1769 | 0 | 0 |
| BMPR2 | 3247 | 0 | 0 |
| PTEN | 1302 | 0 | 0 |
| RPL22 | 619 | 0 | 0 |
| SMAD2 | 1504 | 0 | 0 |
| ATM | 9791 | 0 | 0 |
| POLE | 7351 | 0 | 0 |
| ARID1A | 7058 | 0 | 0 |
| FBXW7 | 2758 | 0 | 0 |
| RNF43 | 2442 | 0 | 0 |
| MLH1 | 2461 | 0 | 0 |
| MSH2 | 3107 | 0 | 0 |
| KRAS | 737 | 0 | 0 |
| ELF3 | 1256 | 0 | 0 |
| POLD1 | 3662 | 0 | 0 |
| CTNNB1 | 2486 | 0 | 0 |
| ZFP36L2 | 1505 | 0 | 0 |
| APC | 8857 | 0 | 0 |
| SOX9 | 1560 | 0 | 0 |
| ACVR2A | 1652 | 0 | 0 |
| chr1 | 6246726 | 6246881 | RPL22 |
| chr1 | 6252984 | 6253119 | RPL22 |
| chr1 | 6257706 | 6257821 | RPL22 |
| chr1 | 6259424 | 6259638 | RPL22 |
| chr1 | 27022889 | 27024036 | ARID1A |
| chr1 | 27056136 | 27056359 | ARID1A |
| chr1 | 27057637 | 27058100 | ARID1A |
| chr1 | 27059161 | 27059288 | ARID1A |
| chr1 | 27087341 | 27087592 | ARID1A |
| chr1 | 27087869 | 27087969 | ARID1A |
| chr1 | 27088637 | 27088815 | ARID1A |
| chr1 | 27089458 | 27089781 | ARID1A |
| chr1 | 27092706 | 27092862 | ARID1A |
| chr1 | 27092942 | 27093062 | ARID1A |
| chr1 | 27094275 | 27094495 | ARID1A |
| chr1 | 27097604 | 27097822 | ARID1A |
| chr1 | 27098985 | 27099128 | ARID1A |
| chr1 | 27099297 | 27099483 | ARID1A |
| chr1 | 27099831 | 27099992 | ARID1A |
| chr1 | 27100065 | 27100213 | ARID1A |
| chr1 | 27100287 | 27100394 | ARID1A |
| chr1 | 27100814 | 27101716 | ARID1A |
| chr1 | 27102062 | 27102203 | ARID1A |
| chr1 | 27105508 | 27107252 | ARID1A |
| chr1 | 115251150 | 115251280 | NRAS |
| chr1 | 115252184 | 115252354 | NRAS |
| chr1 | 115256415 | 115256604 | NRAS |
| chr1 | 115258665 | 115258786 | NRAS |
| chr1 | 201980259 | 201980432 | ELF3 |
| chr1 | 201981079 | 201981311 | ELF3 |
| chr1 | 201981466 | 201981569 | ELF3 |
| chr1 | 201981762 | 201981892 | ELF3 |
| chr1 | 201981973 | 201982033 | ELF3 |
| chr1 | 201982069 | 201982169 | ELF3 |
| chr1 | 201982304 | 201982431 | ELF3 |
| chr1 | 201982951 | 201983157 | ELF3 |
| chr1 | 201984331 | 201984456 | ELF3 |
| chr10 | 89624221 | 89624310 | PTEN |
| chr10 | 89653776 | 89653871 | PTEN |
| chr10 | 89685264 | 89685319 | PTEN |
| chr10 | 89690797 | 89690851 | PTEN |
| chr10 | 89692764 | 89693013 | PTEN |
| chr10 | 89711869 | 89712021 | PTEN |
| chr10 | 89717604 | 89717781 | PTEN |
| chr10 | 89720645 | 89720880 | PTEN |
| chr10 | 89725038 | 89725234 | PTEN |
| chr10 | 114710510 | 114710709 | TCF7L2 |
| chr10 | 114710960 | 114711037 | TCF7L2 |
| chr10 | 114711236 | 114711371 | TCF7L2 |
| chr10 | 114724309 | 114724388 | TCF7L2 |
| chr10 | 114799778 | 114799890 | TCF7L2 |
| chr10 | 114849150 | 114849304 | TCF7L2 |
| chr10 | 114886632 | 114886645 | TCF7L2 |
| chr10 | 114889618 | 114889751 | TCF7L2 |
| chr10 | 114900937 | 114901080 | TCF7L2 |
| chr10 | 114903676 | 114903789 | TCF7L2 |
| chr10 | 114905764 | 114905861 | TCF7L2 |
| chr10 | 114910736 | 114910887 | TCF7L2 |
| chr10 | 114911478 | 114911648 | TCF7L2 |
| chr10 | 114912086 | 114912204 | TCF7L2 |
| chr10 | 114917774 | 114917833 | TCF7L2 |
| chr10 | 114918420 | 114918481 | TCF7L2 |
| chr10 | 114919673 | 114919756 | TCF7L2 |
| chr10 | 114920372 | 114920455 | TCF7L2 |
| chr10 | 114921332 | 114921349 | TCF7L2 |
| chr10 | 114925308 | 114925736 | TCF7L2 |
| chr11 | 108098346 | 108098428 | ATM |
| chr11 | 108098497 | 108098620 | ATM |
| chr11 | 108099899 | 108100055 | ATM |
| chr11 | 108106391 | 108106566 | ATM |
| chr11 | 108114674 | 108114850 | ATM |
| chr11 | 108115509 | 108115758 | ATM |
| chr11 | 108117685 | 108117859 | ATM |
| chr11 | 108119654 | 108119834 | ATM |
| chr11 | 108121422 | 108121804 | ATM |
| chr11 | 108122558 | 108122763 | ATM |
| chr11 | 108123538 | 108123644 | ATM |
| chr11 | 108124535 | 108124771 | ATM |
| chr11 | 108126936 | 108127072 | ATM |
| chr11 | 108128202 | 108128338 | ATM |
| chr11 | 108129707 | 108129807 | ATM |
| chr11 | 108137892 | 108138074 | ATM |
| chr11 | 108139131 | 108139341 | ATM |
| chr11 | 108141785 | 108141878 | ATM |
| chr11 | 108141972 | 108142138 | ATM |
| chr11 | 108143253 | 108143339 | ATM |
| chr11 | 108143443 | 108143584 | ATM |
| chr11 | 108150212 | 108150340 | ATM |
| chr11 | 108151716 | 108151900 | ATM |
| chr11 | 108153431 | 108153611 | ATM |
| chr11 | 108154948 | 108155205 | ATM |
| chr11 | 108158321 | 108158447 | ATM |
| chr11 | 108159698 | 108159835 | ATM |
| chr11 | 108160323 | 108160533 | ATM |
| chr11 | 108163340 | 108163525 | ATM |
| chr11 | 108164034 | 108164209 | ATM |
| chr11 | 108165648 | 108165791 | ATM |
| chr11 | 108168008 | 108168114 | ATM |
| chr11 | 108170435 | 108170617 | ATM |
| chr11 | 108172369 | 108172521 | ATM |
| chr11 | 108173574 | 108173761 | ATM |
| chr11 | 108175396 | 108175584 | ATM |
| chr11 | 108178618 | 108178716 | ATM |
| chr11 | 108180881 | 108181047 | ATM |
| chr11 | 108183132 | 108183230 | ATM |
| chr11 | 108186544 | 108186643 | ATM |
| chr11 | 108186732 | 108186845 | ATM |
| chr11 | 108188094 | 108188253 | ATM |
| chr11 | 108190675 | 108190790 | ATM |
| chr11 | 108192022 | 108192152 | ATM |
| chr11 | 108196031 | 108196276 | ATM |
| chr11 | 108196779 | 108196957 | ATM |
| chr11 | 108198366 | 108198490 | ATM |
| chr11 | 108199742 | 108199970 | ATM |
| chr11 | 108200935 | 108201153 | ATM |
| chr11 | 108202165 | 108202289 | ATM |
| chr11 | 108202600 | 108202769 | ATM |
| chr11 | 108203483 | 108203632 | ATM |
| chr11 | 108204607 | 108204700 | ATM |
| chr11 | 108205690 | 108205841 | ATM |
| chr11 | 108206566 | 108206693 | ATM |
| chr11 | 108213943 | 108214103 | ATM |
| chr11 | 108216464 | 108216640 | ATM |
| chr11 | 108218000 | 108218097 | ATM |
| chr11 | 108224487 | 108224612 | ATM |
| chr11 | 108225532 | 108225606 | ATM |
| chr11 | 108235803 | 108235950 | ATM |
| chr11 | 108236046 | 108236240 | ATM |
| chr11 | 118769118 | 118770222 | BCL9L |
| chr11 | 118770620 | 118770912 | BCL9L |
| chr11 | 118771322 | 118773622 | BCL9L |
| chr11 | 118773693 | 118773788 | BCL9L |
| chr11 | 118773939 | 118774166 | BCL9L |
| chr11 | 118778186 | 118778316 | BCL9L |
| chr11 | 118778973 | 118779369 | BCL9L |
| chr11 | 118780617 | 118780653 | BCL9L |
| chr12 | 25362723 | 25362850 | KRAS |
| chr12 | 25368369 | 25368499 | KRAS |
| chr12 | 25378542 | 25378712 | KRAS |
| chr12 | 25380162 | 25380351 | KRAS |
| chr12 | 25398202 | 25398323 | KRAS |
| chr12 | 133201277 | 133201401 | POLE |
| chr12 | 133201485 | 133201585 | POLE |
| chr12 | 133202225 | 133202361 | POLE |
| chr12 | 133202697 | 133202908 | POLE |
| chr12 | 133208895 | 133209099 | POLE |
| chr12 | 133209244 | 133209386 | POLE |
| chr12 | 133210766 | 133210969 | POLE |
| chr12 | 133212472 | 133212615 | POLE |
| chr12 | 133214594 | 133214730 | POLE |
| chr12 | 133215705 | 133215889 | POLE |
| chr12 | 133218227 | 133218442 | POLE |
| chr12 | 133218757 | 133218988 | POLE |
| chr12 | 133219086 | 133219320 | POLE |
| chr12 | 133219400 | 133219587 | POLE |
| chr12 | 133219804 | 133219921 | POLE |
| chr12 | 133219987 | 133220151 | POLE |
| chr12 | 133220417 | 133220568 | POLE |
| chr12 | 133225509 | 133225663 | POLE |
| chr12 | 133225886 | 133226106 | POLE |
| chr12 | 133226257 | 133226480 | POLE |
| chr12 | 133233716 | 133233849 | POLE |
| chr12 | 133233929 | 133234020 | POLE |
| chr12 | 133234448 | 133234561 | POLE |
| chr12 | 133235875 | 133236100 | POLE |
| chr12 | 133237549 | 133237755 | POLE |
| chr12 | 133238107 | 133238275 | POLE |
| chr12 | 133240584 | 133240739 | POLE |
| chr12 | 133240950 | 133241053 | POLE |
| chr12 | 133241882 | 133242041 | POLE |
| chr12 | 133244083 | 133244239 | POLE |
| chr12 | 133244936 | 133245093 | POLE |
| chr12 | 133245215 | 133245328 | POLE |
| chr12 | 133245391 | 133245530 | POLE |
| chr12 | 133248795 | 133248913 | POLE |
| chr12 | 133249207 | 133249430 | POLE |
| chr12 | 133249744 | 133249868 | POLE |
| chr12 | 133250155 | 133250298 | POLE |
| chr12 | 133251978 | 133252108 | POLE |
| chr12 | 133252315 | 133252411 | POLE |
| chr12 | 133252674 | 133252795 | POLE |
| chr12 | 133253126 | 133253244 | POLE |
| chr12 | 133253943 | 133254034 | POLE |
| chr12 | 133254158 | 133254310 | POLE |
| chr12 | 133256077 | 133256242 | POLE |
| chr12 | 133256534 | 133256637 | POLE |
| chr12 | 133256758 | 133256813 | POLE |
| chr12 | 133257187 | 133257278 | POLE |
| chr12 | 133257718 | 133257870 | POLE |
| chr12 | 133263834 | 133263906 | POLE |
| chr15 | 45003739 | 45003816 | B2M |
| chr15 | 45007615 | 45007927 | B2M |
| chr15 | 45008521 | 45008545 | B2M |
| chr17 | 7572921 | 7573013 | TP53 |
| chr17 | 7573921 | 7574038 | TP53 |
| chr17 | 7576531 | 7576589 | TP53 |
| chr17 | 7576619 | 7576662 | TP53 |
| chr17 | 7576847 | 7576931 | TP53 |
| chr17 | 7577013 | 7577160 | TP53 |
| chr17 | 7577493 | 7577613 | TP53 |
| chr17 | 7578171 | 7578294 | TP53 |
| chr17 | 7578365 | 7578559 | TP53 |
| chr17 | 7579306 | 7579595 | TP53 |
| chr17 | 7579694 | 7579726 | TP53 |
| chr17 | 7579833 | 7579917 | TP53 |
| chr17 | 56432298 | 56432352 | RNF43 |
| chr17 | 56434823 | 56436189 | RNF43 |
| chr17 | 56437504 | 56437617 | RNF43 |
| chr17 | 56438138 | 56438310 | RNF43 |
| chr17 | 56439899 | 56440014 | RNF43 |
| chr17 | 56440630 | 56440772 | RNF43 |
| chr17 | 56440881 | 56440966 | RNF43 |
| chr17 | 56448266 | 56448399 | RNF43 |
| chr17 | 56492681 | 56492943 | RNF43 |
| chr17 | 70117527 | 70117968 | SOX9 |
| chr17 | 70118854 | 70119118 | SOX9 |
| chr17 | 70119678 | 70120533 | SOX9 |
| chr18 | 3447732 | 3447800 | TGIF1 |
| chr18 | 3449624 | 3449659 | TGIF1 |
| chr18 | 3450482 | 3450508 | TGIF1 |
| chr18 | 3451972 | 3452385 | TGIF1 |
| chr18 | 3456346 | 3456583 | TGIF1 |
| chr18 | 3457357 | 3457943 | TGIF1 |
| chr18 | 45368192 | 45368326 | SMAD2 |
| chr18 | 45371705 | 45371860 | SMAD2 |
| chr18 | 45372028 | 45372176 | SMAD2 |
| chr18 | 45374840 | 45375063 | SMAD2 |
| chr18 | 45377639 | 45377703 | SMAD2 |
| chr18 | 45391424 | 45391509 | SMAD2 |
| chr18 | 45394688 | 45394833 | SMAD2 |
| chr18 | 45395608 | 45395812 | SMAD2 |
| chr18 | 45396840 | 45396940 | SMAD2 |
| chr18 | 45422886 | 45423132 | SMAD2 |
| chr18 | 48573411 | 48573670 | SMAD4 |
| chr18 | 48575050 | 48575235 | SMAD4 |
| chr18 | 48575659 | 48575699 | SMAD4 |
| chr18 | 48581145 | 48581368 | SMAD4 |
| chr18 | 48584489 | 48584619 | SMAD4 |
| chr18 | 48584704 | 48584831 | SMAD4 |
| chr18 | 48586230 | 48586291 | SMAD4 |
| chr18 | 48591787 | 48591981 | SMAD4 |
| chr18 | 48593383 | 48593562 | SMAD4 |
| chr18 | 48603002 | 48603151 | SMAD4 |
| chr18 | 48604620 | 48604842 | SMAD4 |
| chr19 | 50902103 | 50902315 | POLD1 |
| chr19 | 50902622 | 50902746 | POLD1 |
| chr19 | 50905029 | 50905186 | POLD1 |
| chr19 | 50905250 | 50905386 | POLD1 |
| chr19 | 50905456 | 50905635 | POLD1 |
| chr19 | 50905705 | 50905797 | POLD1 |
| chr19 | 50905863 | 50906003 | POLD1 |
| chr19 | 50906304 | 50906481 | POLD1 |
| chr19 | 50906744 | 50906859 | POLD1 |
| chr19 | 50909433 | 50909584 | POLD1 |
| chr19 | 50909658 | 50909779 | POLD1 |
| chr19 | 50910234 | 50910436 | POLD1 |
| chr19 | 50910578 | 50910677 | POLD1 |
| chr19 | 50911958 | 50912163 | POLD1 |
| chr19 | 50912373 | 50912497 | POLD1 |
| chr19 | 50912770 | 50912928 | POLD1 |
| chr19 | 50916677 | 50916783 | POLD1 |
| chr19 | 50916993 | 50917141 | POLD1 |
| chr19 | 50918066 | 50918252 | POLD1 |
| chr19 | 50918689 | 50918852 | POLD1 |
| chr19 | 50918975 | 50919088 | POLD1 |
| chr19 | 50919647 | 50919790 | POLD1 |
| chr19 | 50919861 | 50919985 | POLD1 |
| chr19 | 50920296 | 50920359 | POLD1 |
| chr19 | 50920423 | 50920531 | POLD1 |
| chr19 | 50921093 | 50921209 | POLD1 |
| chr2 | 43451452 | 43452896 | ZFP36L2 |
| chr2 | 43453398 | 43453459 | ZFP36L2 |
| chr2 | 47630325 | 47630546 | MSH2 |
| chr2 | 47635534 | 47635699 | MSH2 |
| chr2 | 47637227 | 47637516 | MSH2 |
| chr2 | 47639547 | 47639704 | MSH2 |
| chr2 | 47641402 | 47641562 | MSH2 |
| chr2 | 47643429 | 47643573 | MSH2 |
| chr2 | 47656875 | 47657085 | MSH2 |
| chr2 | 47672681 | 47672801 | MSH2 |
| chr2 | 47690164 | 47690298 | MSH2 |
| chr2 | 47693791 | 47693952 | MSH2 |
| chr2 | 47698098 | 47698206 | MSH2 |
| chr2 | 47702158 | 47702414 | MSH2 |
| chr2 | 47703500 | 47703715 | MSH2 |
| chr2 | 47705405 | 47705663 | MSH2 |
| chr2 | 47707829 | 47708015 | MSH2 |
| chr2 | 47709912 | 47710093 | MSH2 |
| chr2 | 47739436 | 47739578 | MSH2 |
| chr2 | 48010367 | 48010637 | MSH6 |
| chr2 | 48018060 | 48018267 | MSH6 |
| chr2 | 48023027 | 48023207 | MSH6 |
| chr2 | 48025744 | 48028299 | MSH6 |
| chr2 | 48030553 | 48030829 | MSH6 |
| chr2 | 48032043 | 48032171 | MSH6 |
| chr2 | 48032751 | 48032851 | MSH6 |
| chr2 | 48033337 | 48033502 | MSH6 |
| chr2 | 48033585 | 48033795 | MSH6 |
| chr2 | 48033912 | 48034004 | MSH6 |
| chr2 | 148602716 | 148602781 | ACVR2A |
| chr2 | 148653864 | 148654082 | ACVR2A |
| chr2 | 148657021 | 148657141 | ACVR2A |
| chr2 | 148657307 | 148657472 | ACVR2A |
| chr2 | 148672754 | 148672908 | ACVR2A |
| chr2 | 148674846 | 148675000 | ACVR2A |
| chr2 | 148676010 | 148676166 | ACVR2A |
| chr2 | 148677793 | 148677918 | ACVR2A |
| chr2 | 148680536 | 148680685 | ACVR2A |
| chr2 | 148683594 | 148683735 | ACVR2A |
| chr2 | 148684643 | 148684848 | ACVR2A |
| chr2 | 203242192 | 203242278 | BMPR2 |
| chr2 | 203329526 | 203329707 | BMPR2 |
| chr2 | 203332236 | 203332417 | BMPR2 |
| chr2 | 203378436 | 203378557 | BMPR2 |
| chr2 | 203379605 | 203379707 | BMPR2 |
| chr2 | 203383539 | 203383780 | BMPR2 |
| chr2 | 203384804 | 203384929 | BMPR2 |
| chr2 | 203395511 | 203395682 | BMPR2 |
| chr2 | 203397302 | 203397460 | BMPR2 |
| chr2 | 203407028 | 203407175 | BMPR2 |
| chr2 | 203417433 | 203417616 | BMPR2 |
| chr2 | 203419969 | 203421259 | BMPR2 |
| chr2 | 203424413 | 203424674 | BMPR2 |
| chr20 | 57415156 | 57415946 | GNAS |
| chr20 | 57428315 | 57430393 | GNAS |
| chr20 | 57466776 | 57466925 | GNAS |
| chr20 | 57470661 | 57470744 | GNAS |
| chr20 | 57473990 | 57474045 | GNAS |
| chr20 | 57478577 | 57478645 | GNAS |
| chr20 | 57478721 | 57478851 | GNAS |
| chr20 | 57480432 | 57480540 | GNAS |
| chr20 | 57484211 | 57484276 | GNAS |
| chr20 | 57484399 | 57484483 | GNAS |
| chr20 | 57484570 | 57484639 | GNAS |
| chr20 | 57484733 | 57484864 | GNAS |
| chr20 | 57485000 | 57485141 | GNAS |
| chr20 | 57485383 | 57485461 | GNAS |
| chr20 | 57485732 | 57485889 | GNAS |
| chr3 | 37035033 | 37035159 | MLH1 |
| chr3 | 37038104 | 37038205 | MLH1 |
| chr3 | 37042440 | 37042549 | MLH1 |
| chr3 | 37045886 | 37045970 | MLH1 |
| chr3 | 37048476 | 37048559 | MLH1 |
| chr3 | 37050299 | 37050401 | MLH1 |
| chr3 | 37053305 | 37053358 | MLH1 |
| chr3 | 37053496 | 37053595 | MLH1 |
| chr3 | 37055917 | 37056040 | MLH1 |
| chr3 | 37058991 | 37059095 | MLH1 |
| chr3 | 37061795 | 37061959 | MLH1 |
| chr3 | 37067122 | 37067503 | MLH1 |
| chr3 | 37070269 | 37070428 | MLH1 |
| chr3 | 37081671 | 37081790 | MLH1 |
| chr3 | 37083753 | 37083827 | MLH1 |
| chr3 | 37089004 | 37089179 | MLH1 |
| chr3 | 37090002 | 37090105 | MLH1 |
| chr3 | 37090389 | 37090513 | MLH1 |
| chr3 | 37091971 | 37092149 | MLH1 |
| chr3 | 41265554 | 41265577 | CTNNB1 |
| chr3 | 41266011 | 41266249 | CTNNB1 |
| chr3 | 41266439 | 41266703 | CTNNB1 |
| chr3 | 41266819 | 41267068 | CTNNB1 |
| chr3 | 41267145 | 41267357 | CTNNB1 |
| chr3 | 41268693 | 41268848 | CTNNB1 |
| chr3 | 41274826 | 41274940 | CTNNB1 |
| chr3 | 41275014 | 41275363 | CTNNB1 |
| chr3 | 41275624 | 41275793 | CTNNB1 |
| chr3 | 41277209 | 41277339 | CTNNB1 |
| chr3 | 41277834 | 41277995 | CTNNB1 |
| chr3 | 41278073 | 41278205 | CTNNB1 |
| chr3 | 41279501 | 41279572 | CTNNB1 |
| chr3 | 41280619 | 41280838 | CTNNB1 |
| chr3 | 178916608 | 178916970 | PIK3CA |
| chr3 | 178917472 | 178917692 | PIK3CA |
| chr3 | 178919072 | 178919333 | PIK3CA |
| chr3 | 178921326 | 178921582 | PIK3CA |
| chr3 | 178922285 | 178922381 | PIK3CA |
| chr3 | 178927377 | 178927493 | PIK3CA |
| chr3 | 178927968 | 178928131 | PIK3CA |
| chr3 | 178928213 | 178928358 | PIK3CA |
| chr3 | 178935992 | 178936127 | PIK3CA |
| chr3 | 178936978 | 178937070 | PIK3CA |
| chr3 | 178937353 | 178937528 | PIK3CA |
| chr3 | 178937731 | 178937845 | PIK3CA |
| chr3 | 178938768 | 178938950 | PIK3CA |
| chr3 | 178941863 | 178941980 | PIK3CA |
| chr3 | 178942482 | 178942614 | PIK3CA |
| chr3 | 178943744 | 178943833 | PIK3CA |
| chr3 | 178947054 | 178947235 | PIK3CA |
| chr3 | 178947786 | 178947914 | PIK3CA |
| chr3 | 178948007 | 178948169 | PIK3CA |
| chr3 | 178951876 | 178952157 | PIK3CA |
| chr4 | 153244027 | 153244306 | FBXW7 |
| chr4 | 153245330 | 153245551 | FBXW7 |
| chr4 | 153247152 | 153247388 | FBXW7 |
| chr4 | 153249354 | 153249546 | FBXW7 |
| chr4 | 153250818 | 153250942 | FBXW7 |
| chr4 | 153251878 | 153252025 | FBXW7 |
| chr4 | 153253742 | 153253876 | FBXW7 |
| chr4 | 153258948 | 153259093 | FBXW7 |
| chr4 | 153268076 | 153268228 | FBXW7 |
| chr4 | 153269820 | 153269886 | FBXW7 |
| chr4 | 153271188 | 153271281 | FBXW7 |
| chr4 | 153273616 | 153273887 | FBXW7 |
| chr4 | 153303335 | 153303492 | FBXW7 |
| chr4 | 153332419 | 153332960 | FBXW7 |
| chr5 | 112043409 | 112043584 | APC |
| chr5 | 112090582 | 112090727 | APC |
| chr5 | 112102017 | 112102112 | APC |
| chr5 | 112102880 | 112103092 | APC |
| chr5 | 112111320 | 112111439 | APC |
| chr5 | 112116481 | 112116605 | APC |
| chr5 | 112128137 | 112128231 | APC |
| chr5 | 112136970 | 112137085 | APC |
| chr5 | 112151186 | 112151295 | APC |
| chr5 | 112154657 | 112155046 | APC |
| chr5 | 112157587 | 112157693 | APC |
| chr5 | 112162799 | 112162949 | APC |
| chr5 | 112163620 | 112163708 | APC |
| chr5 | 112164547 | 112164674 | APC |
| chr5 | 112170642 | 112170867 | APC |
| chr5 | 112173244 | 112179828 | APC |
| chr7 | 140415822 | 140415841 | BRAF |
| chr7 | 140426161 | 140426321 | BRAF |
| chr7 | 140434391 | 140434575 | BRAF |
| chr7 | 140439606 | 140439751 | BRAF |
| chr7 | 140449081 | 140449223 | BRAF |
| chr7 | 140453069 | 140453198 | BRAF |
| chr7 | 140453981 | 140454038 | BRAF |
| chr7 | 140476706 | 140476893 | BRAF |
| chr7 | 140477785 | 140477880 | BRAF |
| chr7 | 140481370 | 140481498 | BRAF |
| chr7 | 140482815 | 140482962 | BRAF |
| chr7 | 140487342 | 140487389 | BRAF |
| chr7 | 140494102 | 140494272 | BRAF |
| chr7 | 140500156 | 140500286 | BRAF |
| chr7 | 140501206 | 140501365 | BRAF |
| chr7 | 140507754 | 140507867 | BRAF |
| chr7 | 140508686 | 140508800 | BRAF |
| chr7 | 140534403 | 140534677 | BRAF |
| chr7 | 140549905 | 140550017 | BRAF |
| chr7 | 140624360 | 140624508 | BRAF |

**Supplementary Table 2:** PDO mutations identified using the QIASeq™ (Qiagen, Germany) custom targeted DNA sequencing panel

|  | **PDO 411** | **PDO 884** | **PDO 389** | **PDO 653** | **PDO 064** | **PDO 557** |
| --- | --- | --- | --- | --- | --- | --- |
| *APC* | stop gained<br>p.Gln1378* | stop gained<br>p.Tyr1376* | frameshift<br>p.Gln1338fs | frameshift<br>p.Val1479fs | stop gained<br>p.Arg876* | missense<br>p.Ala921Ser |
| *TP53* | missense<br>p.Arg248Gln<br>R248Q | missense<br>p.Arg175His<br>R175H | missense<br>p.Gly245Asp<br>G245D | in frame deletion<br>p.Pro191del<br>P191del | missense<br>p.Arg282Trp<br>R282W | - |
| *MSH6* | frameshift<br>p.Leu1330fs<br>L1330fs | frameshift<br>p.Phe1088fs<br>F1088fs | - | - | - | frameshift<br>p.Phe1104fs<br>F1104fs |
| *SOX9* | - | frameshift<br>p.Tyr297fs<br>Y297fs | frameshift<br>p.Ser323fs<br>S323fs | stop gained<br>p.Gln1378*<br>Q1378* | - | - |
| *TCF7L2* | frameshift<br>p.Lys485fs | - | frameshift<br>p.Lys485fs | - | - | - |
| *PIK3CA* | missense<br>p.Glu545Lys<br>E545K | - | - | - | - | missense<br>p.Arg88Gln<br>R88Q |
| *GNAS* | missense<br>p.Thr225Pro | missense<br>p.Thr225Pro | - | - | - | - |
| *SMAD4* | - | - | missense<br>p.Arg496His | frameshift<br>p.Trp302fs | - | - |
| *BMPR2* | stop gained<br>p.Arg147* | - | - | - | - | frameshift<br>p.Asn583fs |
| *FBXW7* | - | - | - | missense<br>p.Arg465His<br>R465H | missense<br>p.Ser582Leu<br>S582L | - |
| *MSH2* | missense<br>p.Asp167Val | - | frameshift<br>p.Leu458fs<br>L458fs | - | - | - |
| *KRAS* | - | - | - | missense<br>p.Gly12Asp<br>G12D | - | missense<br>p.Gly13Arg<br>G13R |
| *BRAF* | missense<br>p.Val600Glu | - | - | - | - | - |
| *BCL9L* | - | - | - | - | - | frameshift<br>p.Pro449fs |
| *TGIF1* | - | - | - | - | frameshift<br>p.Phe337fs | - |
| *NRAS* | - | missense<br>p.Gly13Arg | - | - | - | - |
| *PTEN* | - | - | - | - | - | missense<br>p.Cys136Tyr |
| *RPL22* | - | - | - | - | - | frameshift<br>p.Lys15fs |
| *POLE* | frameshift<br>p.Val1446fs | - | - | - | - | - |
| *ACVR2A* | - | - | - | - | - | frameshift<br>p.Lys437fs |

**Supplementary Figure 1:**
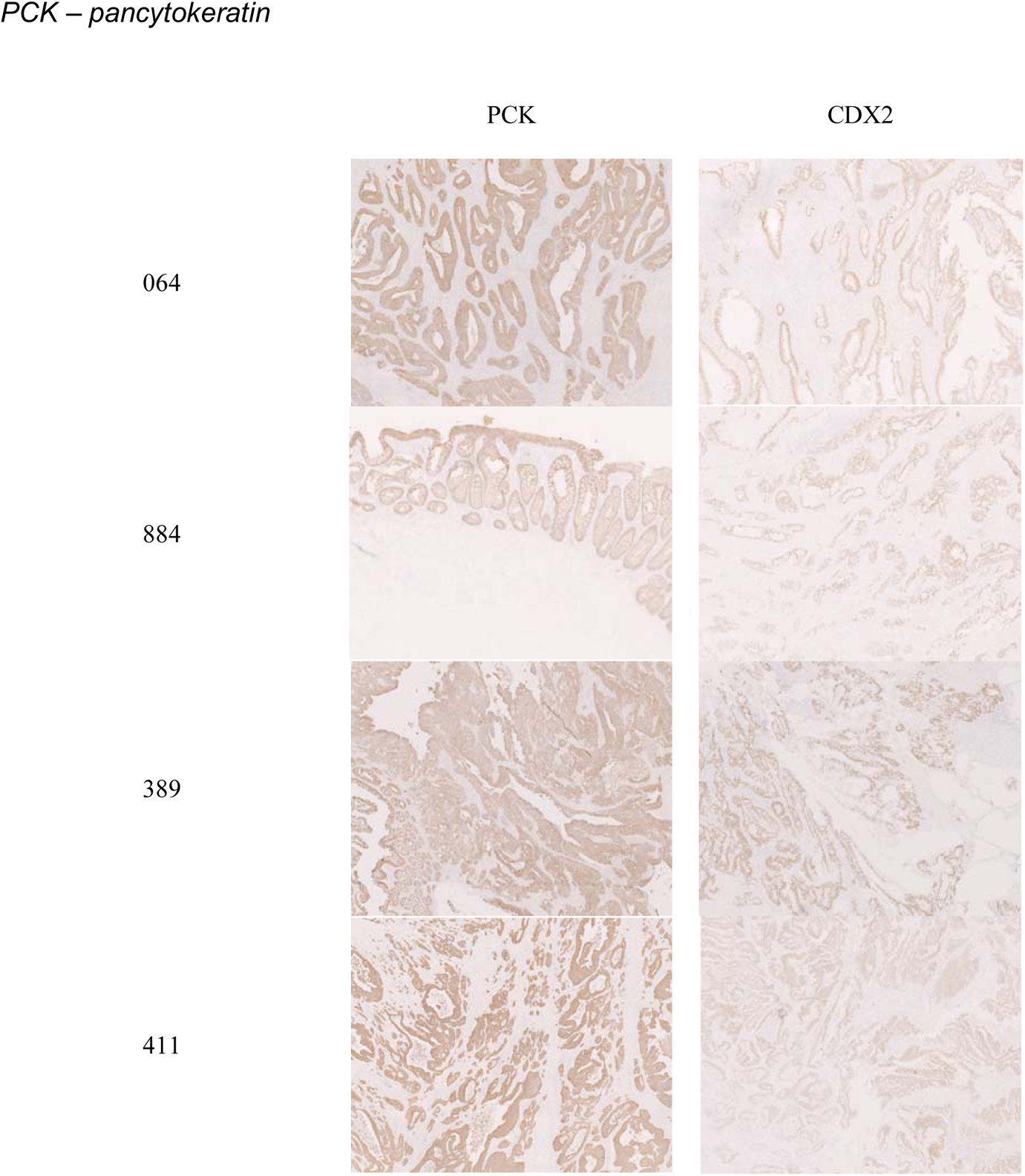
Immunohistochemistry of parent tumour tissue from which PDO lines were derived from.

